# A systems view of spliceosomal assembly and branchpoints with iCLIP

**DOI:** 10.1101/353599

**Authors:** Michael Briese, Nejc Haberman, Christopher R. Sibley, Anob M. Chakrabarti, Zhen Wang, Julian König, David Perera, Vihandha O. Wickramasinghe, Ashok R. Venkitaraman, Nicholas M. Luscombe, Christopher W. Smith, Tomaž Curk, Jernej Ule

## Abstract

Studies of spliceosomal interactions are challenging due to their dynamic nature. Here we employed spliceosome iCLIP, which immunoprecipitates SmB along with snRNPs and auxiliary RNA binding proteins (RBPs), to map human spliceosome engagement with snRNAs and pre-mRNAs. This identified over 50,000 branchpoints (BPs) that have canonical sequence and structural features. Moreover, it revealed 7 binding peaks around BPs and splice sites, each precisely overlapping with binding profiles of specific splicing factors. We show how the binding patterns of these RBPs are affected by the position and strength of BPs. For example, strong or proximally located BPs preferentially bind SF3 rather than U2AF complex. Notably, these effects are partly neutralized during spliceosomal assembly in a way that depends on the core spliceosomal protein PRPF8. These insights exemplify spliceosome iCLIP as a broadly applicable method for transcriptomic studies of splicing mechanisms.

## Introduction

Splicing is a multi-step process in which multiple small nuclear ribonucleoprotein particles (snRNPs) and associated splicing factors bind at specific positions around intron boundaries in order to assemble an active spliceosome through a series of remodeling steps. The splicing reactions are coordinated by dynamic pairings between different snRNAs, between snRNAs and pre-mRNA, and by protein-RNA contacts^1^. Transcriptome-wide studies of splicing reactions can be particularly valuable to unravel the multi-component and dynamic assembly of the spliceosome on the pre-mRNA substrate^2-4^. In yeast, “spliceosome profiling” has been developed through affinity purification of the tagged U2·U5·U6·NTC complex from *Schizosaccharomyces pombe* to monitor its interactions using a RNA footprinting-based strategy^2,3^. It is currently unclear if this method can be applied to mammalian cells, which might be more sensitive to the introduction of affinity tags into splicing factors. Moreover, a method is needed to simultaneously monitor the full complexity of the interactions of diverse RBPs on pre-mRNAs from the earliest to the latest stages of spliceosomal assembly.

A second challenge in understanding splicing mechanisms is the need to assign the position of branchpoints (BPs). The sequence consensus of mammalian BPs is less well defined compared to yeast, therefore experimental methods are important to validate computational predictions. High-throughput methods to identify BPs have so far relied on lariat-spanning RNA-seq reads that cross from the 5’ portion of the intron, over the BP, and finally finish in the 3’ portion of the intron upstream of the BP^5-7^. However, lariat-spanning RNA-seq reads are very rare, and therefore experimental annotation of BPs remains incomplete. In yeast, spliceosome profiling was successful in assigning the positions of BPs by monitoring the position of cDNAs truncating at BPs^2^, indicating that a similar approach could also be applied to mammalian cells.

Here, we have adapted the individual nucleotide resolution UV crosslinking and immunoprecipitation (iCLIP) method^8^ to develop spliceosome iCLIP. This represents a new approach that defines positions of spliceosomal crosslinks on pre-mRNAs at nucleotide resolution^4^ and, thereby, simultaneously maps the crosslink profiles of core and accessory spliceosomal factors that are known to participate across the diverse stages of the splicing cycle. Due to the nucleotide precision of iCLIP, we could distinguish 7 binding peaks, corresponding to distinct RBPs that differ in their requirement for ATP or for the factor PRPF8. Spliceosome iCLIP also purifies intron lariats, which identified BPs in ∼64% of introns within expressed human genes. Compared to the BPs identified by RNA-seq, those identified by spliceosome iCLIP contain more canonical sequence and structural features. We have further examined the binding profiles of spliceosomal RBPs around the BPs. This demonstrates that the assembly of SF3 and associated spliceosomal complexes tends to be determined by a primary BP in most introns, even though alternative BPs are detected by lariat-derived reads. Moreover, we identify complementary roles of U2AF and SF3 complexes in BP definition. Taken together, these findings demonstrate the value of spliceosome iCLIP for transcriptomic studies of BP definition and spliceosomal interactions with pre-mRNAs.

## Results

### Spliceosome iCLIP identifies interactions between splicing factors, snRNAs and pre-mRNAs

SmB/B’ proteins are part of the highly stable Sm core common to all spliceosomal snRNPs except U6^1^, making them suitable candidates for enriching snRNPs via immunopurification. In order to adapt iCLIP for the study of a multi-component machine like the spliceosome, we used antibodies against the endogenous SmB/B’ proteins^9^ using a range of conditions with differing stringency of detergents and salt concentration for the lysis and washing steps (Supplementary Table 1, Fig. 1a and Supplementary Fig. 1a,b). To enable denaturing purification, we generated HEK293 cells stably expressing Flag-tagged SmB and employed urea to purify SmB via a Flag tag, which minimizes co-purification of additional proteins^10^ (‘stringent’ purification, Supplementary Table 1). We observed a 25 kDa band corresponding to the molecular weight of SmB-RNA complex, which was absent in controls (Supplementary Fig. 1c). Next, we employed the standard, non-denaturing iCLIP condition (‘medium’ stringency), which employs a high concentration of detergents in the lysis buffer, and a washing buffer with 1M NaCl (‘medium’ purification, Supplementary Table 1). This disrupts most protein-protein interactions, but can preserve stable complexes such as snRNPs, which is evident by the multiple radioactive bands in addition to the 25 kDa SmB-RNA complex upon treatment with low RNase (Fig. 1b). No radioactive signal was detected if the SmB/B’ antibody was omitted during immunopurification (Fig. 1b and Supplementary Fig. 1d). To co-purify additional accessory splicing factors, we further decreased the concentration of detergents in the lysis buffer, and used only 0.1M NaCl in the washing buffer (‘mild’ purification, Supplementary Table 1). Under this condition, the diffuse signal at 30-200 kDa strongly increased compared to the medium condition, indicating that the mild condition allows the most efficient purification of proteins associated with snRNPs (Fig. 1a and Supplementary Fig. 1e). Under the low RNase treatment, snRNAs remain more intact, and they can thereby serve as a scaffold for purifying the multi-protein spliceosomal complex (Fig. 1a). A similar radioactive labeling pattern was obtained when using three different monoclonal SmB/B’ antibodies (Supplementary Fig. 1d).

**Fig. 1:**
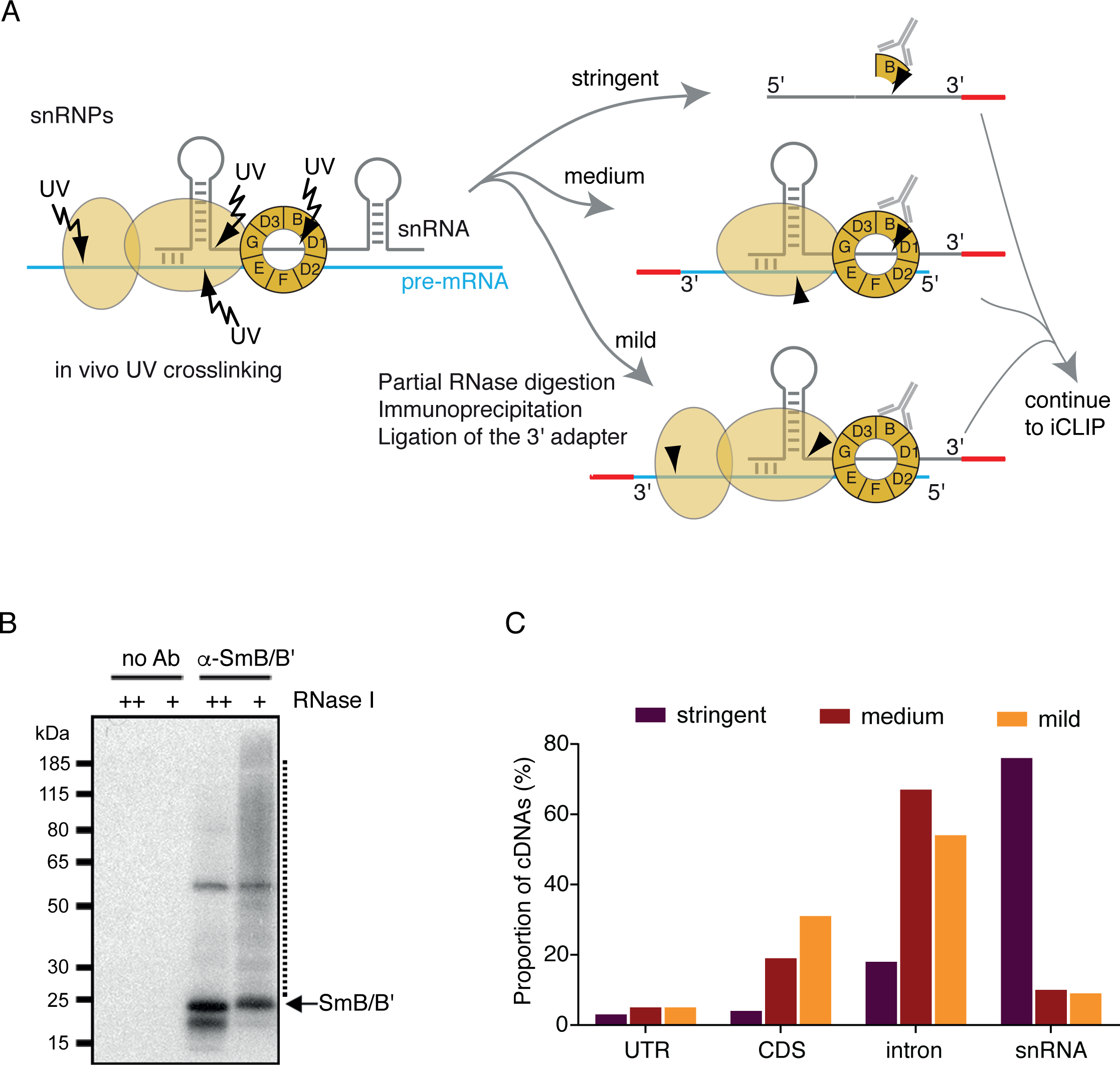
Spliceosome iCLIP identifies protein interactions with snRNAs and splicing substrates. (a) Schematic representation of the spliceosome iCLIP method performed under conditions of varying purification stringency. (b) Autoradiogram of crosslinked RNPs immunopurified from HeLa cells under medium conditions by a SmB/B’ antibody following digestion with high (++) or low (+) amounts of RNase I. The dotted line depicts the region typically excised from the nitrocellulose membrane for spliceosome iCLIP. As control, the antibody (Ab) was omitted during immunopurification. (c) Genomic distribution of spliceosome iCLIP cDNAs. For the analysis cDNAs mapping to untranslated regions (UTR), coding sequence (CDS), introns and snRNAs were considered. For spliceosome iCLIP under medium and mild conditions mouse brain tissue was used and spliceosome iCLIP under stringent conditions was performed on HEK293 cells stably expressing Flag-tagged SmB.

To produce cDNA libraries with spliceosome iCLIP, we immunoprecipitated SmB under the three different stringency conditions from lysates of UV-crosslinked cells or tissue, and isolated a broad size distribution of protein-RNA complexes in order to recover the greatest possible diversity of spliceosomal protein-RNA interactions (Fig. 1b and Supplementary Fig. 1c-e). Mouse brain tissue was used for medium and mild purification with an antibody against endogenous SmB/B’, and HEK293 cells expressing Flag-tagged SmB for stringent, denaturing purification with anti-Flag antibody. As in previous iCLIP studies^8^, the nucleotide preceding each cDNA was used for all analyses. When stringent conditions were used, >75% of iCLIP cDNAs mapped to snRNAs, likely corresponding to the direct binding of Flag-tagged SmB (Fig. 1c). However, the proportion of snRNA crosslinking was reduced to approximately 10% under mild and medium conditions, with a corresponding increase of crosslinking to introns and exons, which likely reflects binding of snRNP-associated proteins to pre-mRNAs (Fig. 1a,c).

### Spliceosome iCLIP identifies seven crosslinking peaks on pre-mRNAs

Assembly of the spliceosome on pre-mRNA is guided by three main landmarks: the 5’ss, 3’ss and BP. Therefore, we evaluated if spliceosomal crosslinks are located at specific positions relative to boundaries of annotated exons and to computationally predicted BPs^11^. For this purpose, we performed spliceosome iCLIP from human Cal51 cells, which have been use as a model system to study the roles of spliceosomal factors in cell cycle^4^. RNA maps of summarized spliceosomal crosslinking revealed 7 peaks of crosslinking around these landmarks, with same positional pattern in Cal51 cells and mouse brain (Fig. 2a and Supplementary Fig. 2a). The centers of the peaks were seen at 15 nt upstream of the 5’ss (peak 1), 10 nt downstream of the 5’ss (peak 2), 31 nt downstream of the 5’ss (peak 3), 26 nt upstream of the BP (peak 4), 20 nt upstream of the BP (peak 5), 11 nt upstream of the 3’ss (peak 6) and 3 nt upstream of the 3’ss (peak 7). We also observed alignment of cDNA starts to the start of the intron and the BPs, which we refer to as positions A and B which are discussed below in more detail (Fig. 2a and Supplementary Fig. 2a).

**Fig. 2:**
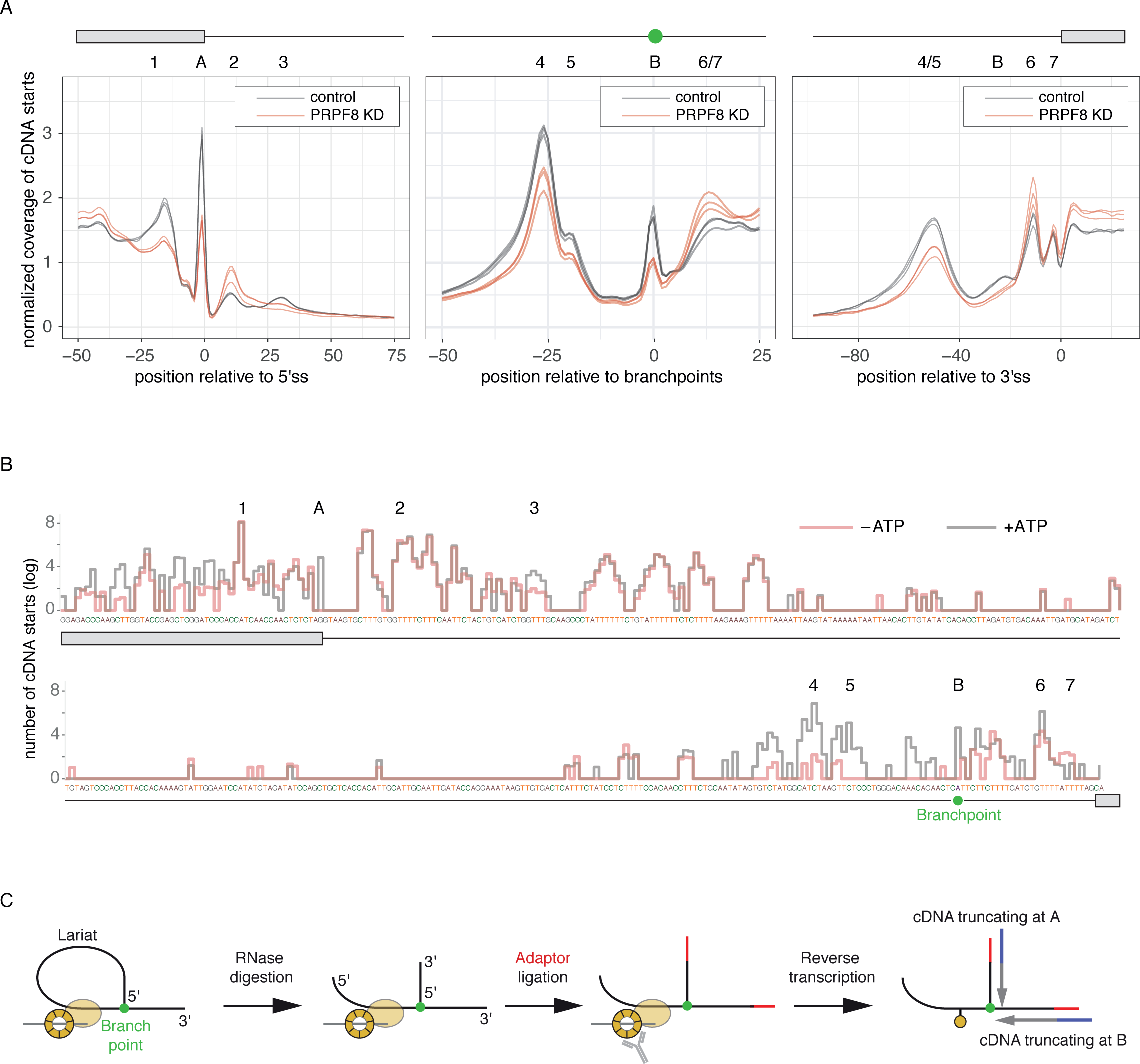
Analysis of spliceosomal interactions with pre-mRNAs *in vitro* and *in vivo*. (a) Metagene plots of spliceosome iCLIP from Cal51 cells. Plots are depicted as RNA maps of summarized crosslinking at all exon-intron and intron-exon boundaries, and around BPs to identify major binding peaks, and to monitor changes between control and PRPF8 knockdown (KD) cells. Crosslinking is regionally normalized to its average crosslinking across the −100..50 nt region relative to 3’ss in order to focus the comparison on the relative positions of peaks. (b) Spliceosome iCLIP cDNA counts on the *C6orf10 in vitro* splicing substrate. Exons are marked by grey boxes, intron by a line, and the BP by a green dot. The positions of crosslinking peaks are marked by numbers and letters corresponding to the peaks in Figure 2a. (c) Schematic description of the three-way junctions of intron lariats. The three-way junction is produced after limited RNase I digestion of intron lariats. This can lead to cDNAs that don’t truncate at sites of protein-RNA crosslinking, but rather at the three-way junction of intron lariats. These cDNAs initiate from the end of the intron and truncate at the BP (position B), or initiate downstream of the 5’ splice site and truncate at the first nucleotide of the intron (position A).

The enrichment of crosslinking at most peaks was generally stronger under the mild condition, especially at the 3’ss, in agreement with the stronger signal of co-purified complexes on the SDS-PAGE gel (Supplementary Fig. 1e and 2a). This indicates that spliceosome iCLIP performed under mild conditions is most suitable for investigating spliceosomal assembly on pre-mRNAs. We therefore used the mild condition to investigate how PRPF8 knockdown (KD) affects spliceosomal interactions in Cal51 cells (Supplementary Fig. 2b). PRPF8 is an integral U5 snRNP component, and therefore part of complexes B and C, where it contacts residues of U5 and U6 snRNAs, as well as pre-mRNA at both the splice sites and BP^1^. We have previously used spliceosome iCLIP to show that PRPF8 is essential for efficient spliceosomal assembly at 5’ss^4^. Here we additionally find that PRPF8 is essential for efficient spliceosomal assembly at peaks 4-5 (Fig. 2a). Moreover, we also observed a major decrease of reads truncating at the positions A and B, whereas crosslinking at peaks 2 and 6 is increased upon PRPF8 KD. Thus, the peaks of spliceosomal crosslinking vary in their sensitivities to PRPF8 depletion.

### *In vitro* spliceosome iCLIP defines the ATP-dependence of crosslinking peaks

In order to verify that spliceosome iCLIP is able to represent multiple stages of the splicing reaction, we performed an *in vitro* splicing assay using defined conditions. We added an exogenous pre-mRNA splicing substrate to HeLa nuclear extract in the presence or absence of ATP. The RNA substrate was produced by *in vitro* transcription of a minigene construct containing a short intron and flanking exons from the human *C6orf10* gene. Gel electrophoresis analysis of splicing products confirmed that ATP was required for the formation of intron lariats and other splicing products (Supplementary Fig. 2c). We performed spliceosome iCLIP from the splicing reactions using the mild purification condition (Supplementary Fig. 2d). Upon sequencing, the reads mapping to the exogenous splicing substrate or the spliced product represented ∼1%, whereas the remaining 99% of mapped reads were derived from endogenous RNAs that are present in the nuclear extract. The spliced product was detected with exon-exon junction reads primarily in the presence of ATP (364 reads in +ATP vs. 5 reads in -ATP condition) (Supplementary Fig. 2e and Supplementary Table 4). Of note, in the +ATP condition the reads mapping to the spliced product (364 reads) were much lower compared to those mapping to the unspliced substrate (48,584 reads) (Supplementary Table 4), as expected given that the spliceosome rapidly disassembles upon completion of the splicing reaction.

We visualized the crosslinking on the substrate RNA, and marked the positions of peaks that corresponded best to those found on endogenous transcripts (Fig. 2b). Whilst crosslinking sites detected on a metagene plot might not necessarily be representative of individual splicing substrates, we nevertheless observed crosslinking peaks in regions of the *C6orf10* substrate at similar positions to the transcriptome-wide peaks (comparing Fig. 2a and 2b). When comparing crosslinking in the presence or absence of ATP, a reproducible crosslinking profile was seen at peaks 1, 2, 6 and 7, indicating that these crosslinks correspond to ATP-independent contacts of early spliceosomal factors. In contrast, the presence of ATP increased the signal at several other peaks: we observed a ∼9 fold increase at peaks 4 and 5, located upstream of the BP, which are also dependent on PRPF8 *in vivo* (Fig. 2a). This indicates that spliceosome iCLIP detects pre-mRNA binding of factors that contribute to distinct stages of spliceosomal assembly.

### Lariat-derived reads are readily obtained by spliceosome iCLIP

Following crosslinking, the peptide that remains bound to the RNA after digestion of the RBP can lead to termination of reverse transcription and produce the so-called ‘truncated cDNAs’^12^. The predominance of truncated cDNAs in iCLIP libraries has been validated by multiple means^13,14^, and therefore our analysis of iCLIP data generally refers to the nucleotide preceding the iCLIP read on the reference genome as the ‘crosslink site’. The same applies to derived methods, such as eCLIP^15^. In spliceosome iCLIP, we expect that cDNAs could also truncate at the three-way junction formed by intron lariats, where the 5’ end of the intron is linked via a 2’-5’ phosphodiester bond to the BP (Fig. 2c). Such three-way-junction RNAs present two available 3’ ends for ligation to adapters, and these reads could truncate at the BP (i.e. position B) or at the start of the intron (i.e. position A), especially if the RBP crosslink site is located upstream of the BP. Indeed, we find strong alignment of cDNA starts at positions A and B, which is dramatically decreased under conditions that decrease the presence of intron lariats: PRPF8 KD *in vivo* (2-fold, Fig. 2a), or the absence of ATP *in vitro* (>15-fold, Fig. 2b). Interestingly, the medium purification condition was optimal to produce cDNAs that truncate at the positions A and B (Supplementary Fig. 2a), possibly because spliceosomal C complexes are readily obtained under high-salt conditions^16^.

### Spliceosome iCLIP identifies >50,000 human branchpoints (BPs)

We performed twelve spliceosome iCLIP experiments under medium purification conditions from UV-crosslinked Cal51 cells that were synchronized at 4 stages of cell cycle, with three replicates for each stage (see Methods). We first confirmed that the starts of spliceosome iCLIP cDNAs generally overlap with a uridine-rich motif (Fig. 3a), in agreement with the increased propensity of protein-RNA crosslinking at uridine-rich sites^13^. In contrast, the nucleotide composition at the starts of cDNAs that end at the last nucleotide of introns strongly overlaps with the YUNAY motif, the consensus sequence of BPs (Fig. 3b). Further, these cDNAs have higher enrichment of mismatches of adenosines at their first nucleotide (Supplementary Fig. 3a), which is consistent with mismatch, insertion and deletion errors during reverse transcription across the three-way junction of the BP^7^. Thus, cDNAs overlapping with intron ends appear to be derived from intron lariats, such that they truncate at the three-way junctions at BPs rather than at crosslink sites of RBPs. In total, they identify 132,287 sites in introns, which could be considered as candidates for BP positions (Fig. 3b).

**Fig. 3:**
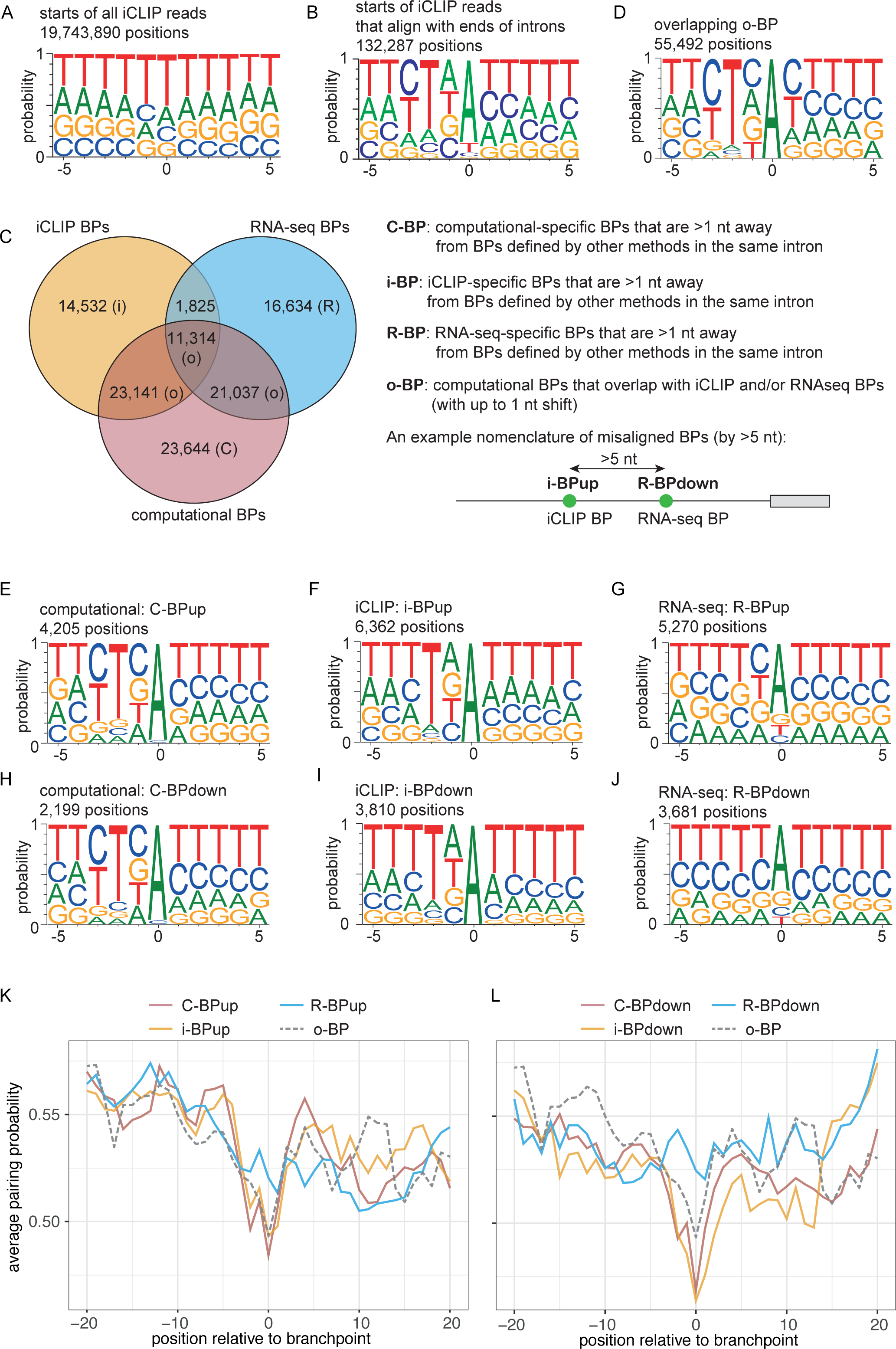
Comparison of BPs identified by spliceosome iCLIP, RNA-seq lariat reads or computational prediction. (a) Weblogo around the nucleotide preceding all spliceosome iCLIP reads. (b) Weblogo around the nucleotide preceding only those spliceosome iCLIP reads that align with ends of introns. (c) Introns that contain at least one BP identified either by published RNA-seq^6^ or by spliceosome iCLIP are used to examine the overlap between the top BPs identified by RNA-seq (i.e., the BP with most lariat-spanning reads in each intron), iCLIP (BP with most cDNA starts) or computational predictions (highest scoring BP)^11^. BPs that are 0 or 1 nt apart are considered as overlapping. At the right, the explanation is given of the BP categories that are used for all subsequent analyses, along with their acronyms. If a BP defined by one method is >5 nt upstream of a BP defined by another method, then ‘up’ is added to its acronym, and if it’s >5 nt downstream, ‘down’ is added. (d) Weblogo of o-BP category of BPs. (e) Weblogo of C-BPup category of BPs. (f) Weblogo of i-BPup category of BPs. (g) Weblogo of R-BPup category of BPs. (h) Weblogo of C-BPdown category of BPs. (i) Weblogo of i-BPdown category of BPs. (j) Weblogo of R-BPdown category of BPs. (k, l) The 100 nt RNA region centered on the BP was used to calculate pairing probability with RNAfold program with the default parameters^28^, and the average pairing probability of each nucleotide around BPs is shown for the 40 nt region around method-specific BPs located upstream (k) or downstream (l).

To identify a confident set of putative BPs in a transcriptome-wide manner, we used the spliceosome iCLIP cDNAs that overlap with intron ends in 9,363 genes with FPKM>10 (as determined by RNA-seq) in Cal51 cells. Thereby we wished to ensure that the genes were expressed at a level that was sufficient for confident analysis of introns. Initially, we only used those cDNAs that overlapped with the end of introns, since we found that these cDNAs tend to start at a BP consensus motif (Fig. 3b). These cDNAs started at adenines in 35,056 introns, which we considered as putative BPs. The more distal BPs would not be identified by this approach due to our 41 read-length limit, and therefore we proceeded to a second step in introns where the initial approach did not identify any BPs. We analyzed all cDNAs, and overlapped their truncation sites with BPs computationally predicted in 2010^17^, in order to maintain independence from the more recently computationally predicted BPs that are used for later comparisons in our paper^11^. We selected the positions of computationally predicted BPs with the highest number of truncated cDNAs, which identified candidate BPs in another 15,756 introns. Collectively, this identified candidate BPs in 50,812 introns of 9,363 genes. These genes in total contain 78,894 annotated introns, and thus iCLIP identified putative BPs in 64% of introns in expressed genes.

### BPs identified by iCLIP contain canonical sequence and structural features

To examine the 50,812 BPs identified by spliceosome iCLIP (‘iCLIP BPs’), we compared them with the ‘computational BPs’ identified recently with a sequence-based deep learning predictor, LaBranchoR, which predicted a BP for over 90% of 3’ss^11^. We also compared with the ‘RNA-seq BPs’, including the 130,294 BPs from 50,810 introns that were identified by analysis of lariat-spanning reads from 17,164 RNA-seq datasets^6^. 61% of iCLIP BPs overlapped with the top-scoring computational BPs (Supplementary Fig. 3b). Interestingly, in cases where iCLIP and computational BPs are located <5 nt apart, they tend to occur within A-rich sequences (Supplementary Fig. 3c). This mismatch could be of technical nature, as truncation of iCLIP cDNAs may not be always precisely aligned to the BPs in case of A-rich sequences, or alternatively multiple As might be capable of serving as BPs when they are located in close vicinity. We therefore allowed 1 nt shift for comparison between methods, as has been done previously^11^, which showed that 68% of iCLIP BPs overlapped with the top-scoring computational BPs, and 26% overlapped with the RNA-seq BPs (Fig. 3c). If the computational BPs overlapped either with an iCLIP BP and/or RNA-seq BP, it generally had a strong BP consensus motif (o-BP, Fig. 3d).

To gain insight into the features of BPs that are unique to each method, we then focused on BPs that were identified by a single method and were >5 nt away from BPs identified by other methods. Notably, the computational-or iCLIP-specific BPs have a strong enrichment of the consensus YUNAY motif (c-BP, i-BP, Fig. 3e,f,h,i). In contrast, the RNA-seq-specific BPs contain a larger proportion of non-canonical BP motifs, which agrees with previous observations^5,7,11^ (Fig. 3g,j). To evaluate this further, we compared the iCLIP BPs with two studies that identified 59,359 BPs by exoribonuclease digestion and targeted RNA-sequencing^7^, and 36,078 BPs by lariat-spanning reads refined by U2snRNP/pre-mRNA base-pairing models^5^. Considering the introns that contained BPs defined both by RNA-seq and iCLIP, we found 55% and 45% overlapping BPs to each study (Supplementary Fig. 3d-g). Again, the iCLIP-specific BPs were more strongly enriched in the consensus YUNAY motif compared the BPs that are specifically identified by either RNA-seq method (Supplementary Fig. 3h-m).

Finally, we examined the local RNA structure around each category of BPs. Overlapping, iCLIP-specific and computational-specific BPs had a strong propensity for single-stranded RNA at the position of the BP, which was not seen for the RNA-seq-specific BPs (Fig. 3k,l). This indicates that the RNA-seq-specific BPs might be structurally less accessible for pairing with U2 snRNP. In conclusion, we find that BPs identified by spliceosome iCLIP contain the expected sequence and structural features.

### Specific RBPs are enriched at each peak of spliceosomal crosslinking

Next, we assessed which RBPs might correspond to the peaks identified by spliceosome iCLIP to play a role in BP recognition (peaks 4-7) or formation of intron lariats (positions A and B). We examined published iCLIP data produced in our lab for 18 previously studied RBPs^18-22^, and eCLIP data from K562 and HepG2 cells for 110 RBPs provided by the ENCODE consortium^15^ to assess normalized crosslinking at each peak. This identified a set of RBPs enriched at each peak (Fig. 4 and Supplementary Table 5). As expected, SF3 components SF3B4, SF3A3 and SF3B1 bind to peaks 4-5^23^, U2AF2 binds the polypyrimidine (polyY) tract (peak 6), and U2AF1 close to the intron-exon junction (peak 7)^21^.

**Fig. 4:**
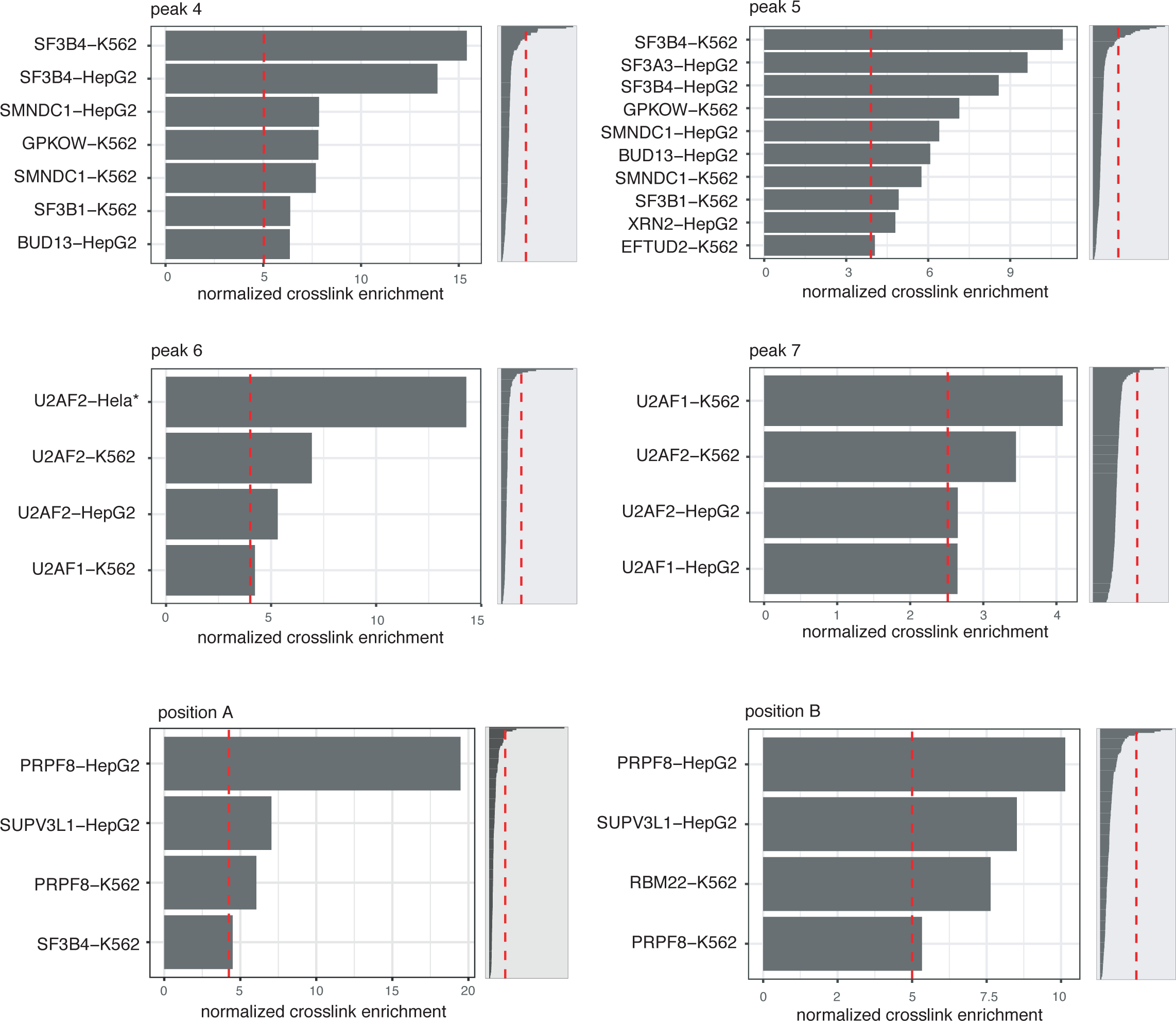
Identification of RBPs overlapping with spliceosomal peaks at BPs and 3’ss. To systematically identify RBPs with crosslinking peaks that overlap with each of the peaks in spliceosome iCLIP, we first regionally normalized the crosslinking of each RBP to its average crosslinking over −100..50 nt region relative to 3’ss, to generate the RNA maps for each RBP as shown in Supplementary Fig. 5 and 6. We then ranked the RBPs according to the the average normalized crosslinking across the nucleotides within each peak. We analyzed peaks 4-7 and positions A and B, as marked on the top of each plot. The top-ranking RBPs in each peak are shown on the left plot, and the full distribution of RBP enrichments is shown on the right plot.

### RBP binding profiles signify the functionality of BPs

Peaks 4-6 and position B align to BP position, and therefore we could evaluate how the crosslinking profiles of RBPs that bind at these peaks align to the different classes of BPs. First, we examined the crosslinking of SF3B4, which binds in the region of peak 4 as part of the U2 snRNP complex that recognises the BP^1^. Analysis of the overlapping BPs (o-BP) defines the peak of SF3B4 crosslinking at the 25^th^ nt upstream of BPs (Fig. 5 and Supplementary Fig. 4a,b). However, the peak of SF3B4 crosslinking doesn’t overlap as well to this 25^th^ position for the non-overlapping, method-specific BPs; it is generally closer than 25 nt to the BPs that are located upstream of another BP (up BP), and further than 25 nt awat from BPs that are located downstream of another BP (down BP) (Fig. 5). The shift from the expected position is greatest for the RNA-seq-specific BPs (R-BP), and smallest for the computationally predicted BPs, as evident by eCLIP data from two cell lines (Fig. 5a,b). Moreover, the same result is seen with U2AF2, where the strongest shift away from expected positions is seen for RNA-seq BPs, and weakest for computational BPs (Supplementary Fig. 4c,d). Given that computationally predicted BPs align best to the SF3 and U2AF binding profiles, we conclude that spliceosome assembles most efficiently on these BPs.

**Fig. 5:**
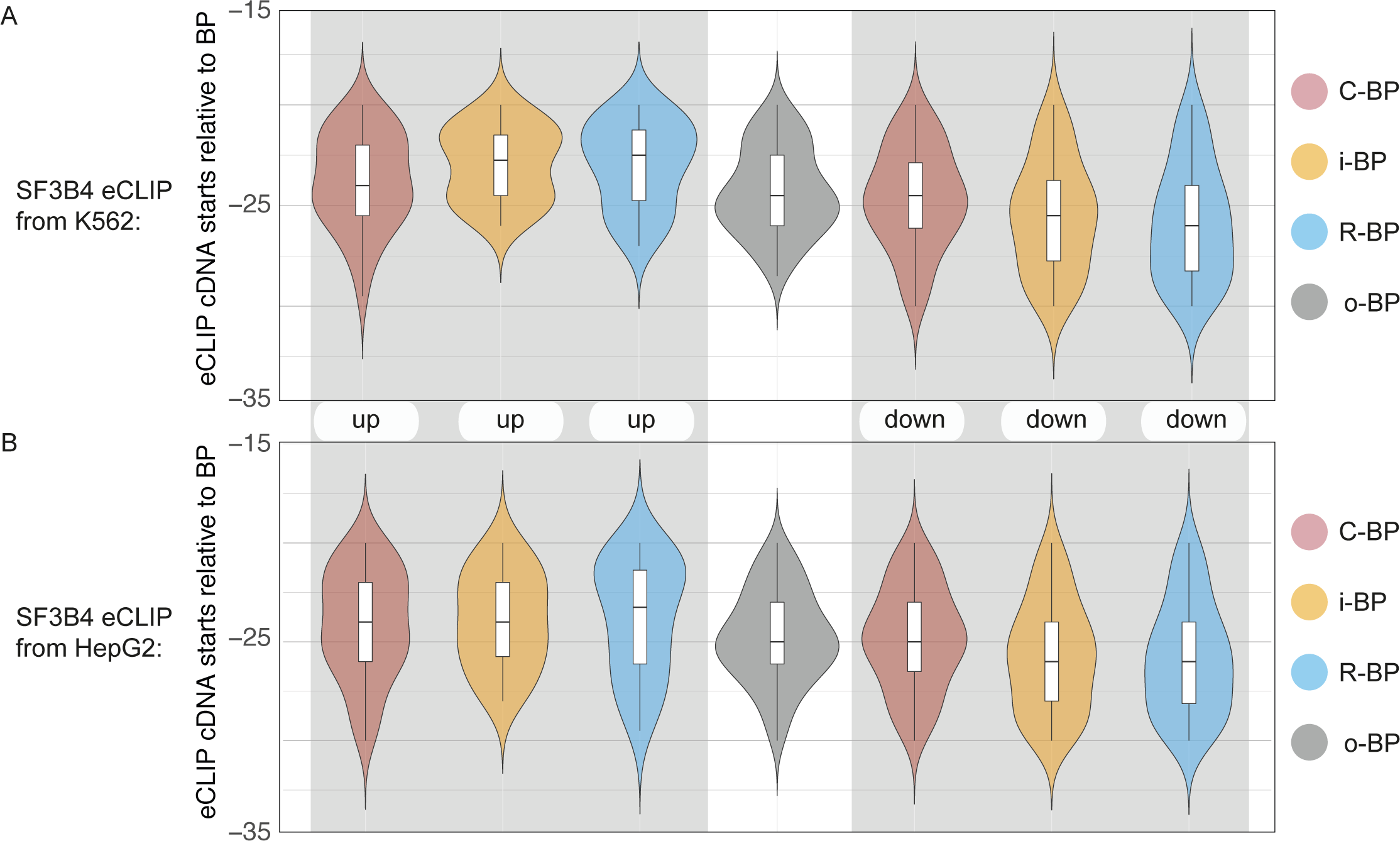
Spliceosome assembly at BPs identified by spliceosome iCLIP, RNA-seq lariat reads or computational prediction. Violin plots depicting the positioning of SF3B4 cDNA starts relative to the indicated BP categories. SF3B4 eCLIP data were from K562 (a) and HepG2 (b) cells. Box-plot elements are defined by center line, median; box limits, upper and lower quartiles; and whiskers, 1.5x interquartile range.

The cDNA starts from PRPF8 eCLIP are highly enriched at position B, corresponding the lariat-derived cDNAs that truncate at BPs (Fig. 4). Interestingly, the PRPF8 cDNA starts had the strongest peak at the overlapping BPs, but also peaked at all the remaining classes of BPs (Supplementary Fig. 4e,f). This indicates that all classes of BPs contribute to lariat formation, and thus the non-overlapping BPs most likely act as alternative BPs within the introns.

### Effects of branchpoint position on spliceosomal assembly

To assess how the position of BPs determines spliceosome assembly, we evaluated the binding profiles of the RBPs that are enriched at peaks 4-7 and at positions A and B (Fig. 4). We divided BPs based on their distance from 3’ss, and normalized the RBP binding profiles within each subclass of BPs. This showed that crosslinking of U2AF1 and U2AF2 aligns to the region between the BPs and 3’ss, which is covered by the polyY tract (Supplementary Fig. 5 and 6). SF3B4 is the primary RBP crosslinking at peak 4, and SF3A3 at 5, and SMNDC1, SF3B1, EFTUD2, BUD13, GPKOW and XRN2 bind to peaks 4/5 (Supplementary Fig. 5, 6 and Fig. 4). PRPF8, RBM22 and SUPV3L1 have their cDNA starts truncating at positions A and B (Supplementary Fig. 5 and 6), corresponding to the three-way junction formed by intron lariats (Fig. 2c), in agreement with the association of PRPF8 and RBM22 with intron lariats as part of the human catalytic step I spliceosome^1^.

In order to quantify how the position of BPs affects the intensity of RBP binding, we divided BPs into 10 equally sized groups based on the distance from 3’ss. We then normalized the relative binding intensity of each RBP at each position on the RNA maps across the ten groups, which revealed strong relationships between BP position and binding intensity of certain RBPs (Fig. 6a, Supplementary Fig. 7a). For example, if a BP is located distally from the 3’ss, then U2AF components bind stronger to peaks 6/7. In contrast, if a BP is located proximally to the 3’ss, then EFTUD2, SF3 components and several other RBPs bind stronger to the peaks 4 or 5 (Fig. 6b). Notably, increased BP distance causes increased binding of BUD13 and GPKOW at peaks 6/7 and decreased binding at peaks 4/5. The more efficient recruitment of U2AF and associated factors to peaks 6/7 could be explained by the long polyY-tracts at distal BPs (Supplementary Fig. 5), while their decreased binding at proximal BPs appears to be compensated for by the increased binding of SF3 and other U2 snRNP-associated factors at peaks 4/5.

**Fig. 6:**
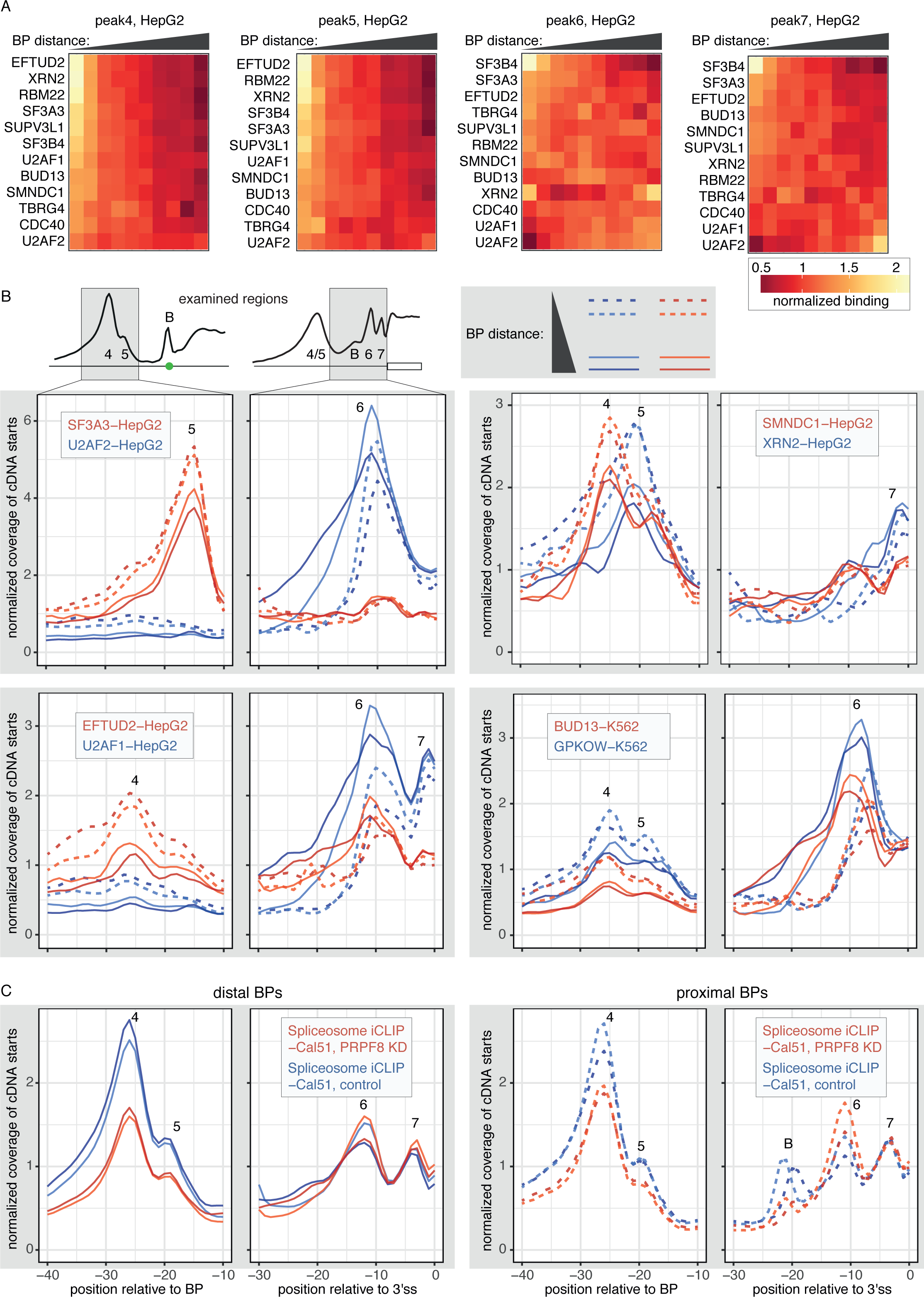
BP position defines the binding patterns of splicing factors at 3’ss. (a) Heatmaps depicting the normalized crosslinking of RBPs in peak regions around 10 groups of BPs that were categorized according to the distance of the BP from 3’ss. Crosslinks were derived as cDNA starts from eCLIP of HepG2 cells. (b) RNA maps showing normalized crosslinking profiles of selected RBPs relative to BPs and 3’ss the two deciles of BPs that are located most proximal (interrupted lines) or most distal (solid lines) from 3’ss. (c) RNA maps showing crosslinking profile of spliceosome iCLIP from control and PRPF8 KD Cal51 cells in the same format as panel b.

In contrast to the effects on individual splicing factors, the relative intensity of spliceosome iCLIP crosslinking in peaks 4/5 compared to 6/7 was not visibly changed in relation to BP distance (Fig. 6c). This indicates that the differences in the binding patterns of individual splicing factors might be neutralized during spliceosomal assembly. To ask if this is the case, we turned to PRPF8, a protein that is essential for the last stage of spliceosome assembly, a role it plays together with EFTUD2 and BRR2 as part of U5 snRNP^1^. PRPF8 knockdown leads to decreased spliceosomal binding at peaks 4/5, and this effect is stronger at distal compared to proximal BPs (Fig. 6c). In conclusion, our results reveal differences in the binding profiles of splicing factors in relation to BP distance, but these differences are neutralized upon spliceosome assembly in a manner that requires the presence of PRPF8.

### Effects of branchpoint strength on spliceosomal assembly

We also wished to examine how the strength of consensus BP sequence affects spliceosomal assembly. For this purpose, we focused on BPs that are located at 23-28 nt upstream of the 3’ss, which is the most common positions of BPs (20,018 BPs, Supplementary Table 6). As an estimate of BP strength we used the BP score, which was determined with a deep-learning model^11^. This showed strong correlation between BP strength and binding intensity of certain RBPs (Fig. 7a, Supplementary Fig. 7b). Among others, increased binding of U2AF is seen at peak 7 of weak BPs, and increased binding of SF3B4 at peaks 4/5 of strong BPs (Fig. 7b). Notably, an over 4-fold change is seen in the ratio between the U2AF and SF3 complexes when comparing the extreme deciles of BP strength (p<0.001, Wilcoxon Rank Sum test, Supplementary Fig. 7c). We did not observe any correlation between the polyY tract coverage and BP score, which indicate that the change in binding profiles is a direct result of BP consensus variation (Supplementary Fig. 7d). Notably, in case of several RBPs, such as XRN2 and SF3B1, weak BP scores correlated with a strong decrease in binding at peaks 6/7 as well as an increase in binding at peaks 4/5 (Fig. 7b).

**Fig. 7:**
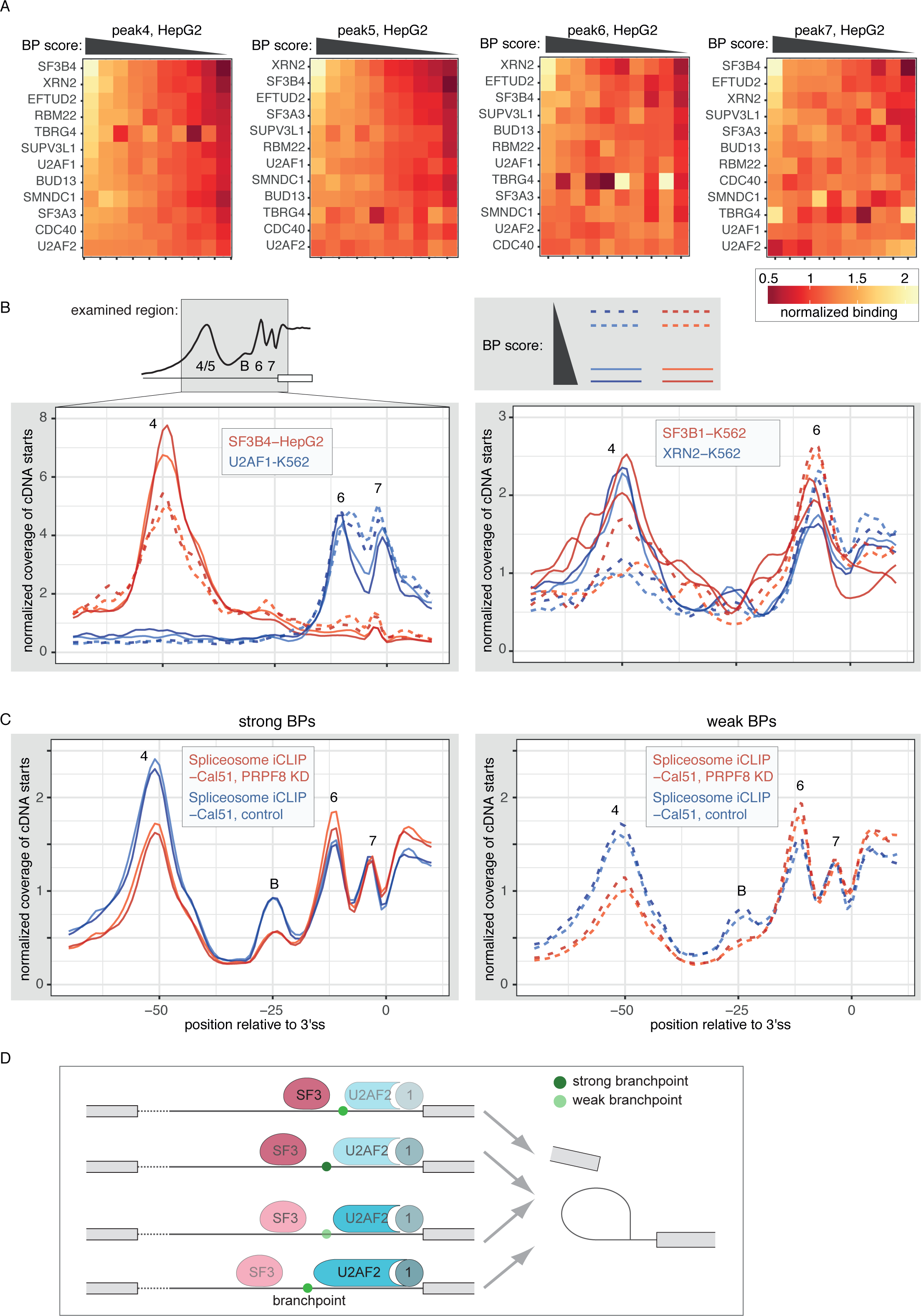
RNA structure around BPs correlates with the binding of splicing factors. (a) Heatmaps depicting the normalized crosslinking of RBPs in peak regions around 10 groups of BPs that were categorized according to the computational scores that define BP strength. Crosslinks were derived as cDNA starts from eCLIP of HepG2 cells. (b) RNA maps showing normalized crosslinking profiles of selected RBPs relative to BPs and 3’ss the two deciles of BPs that are lowest scoring (interrupted lines) or highest scoring (solid lines). (c) RNA maps showing crosslinking profile of spliceosome iCLIP from control and PRPF8 KD Cal51 cells in the same format as panel b. (d) Schematic representation of the effects that BP position and score have on the assembly of SF3 and U2AF complexes around BPs.

Similar to the effects on individual splicing factors, the relative intensity of spliceosome iCLIP crosslinking in peaks 4/5 was increased with increasing BP strength (Fig. 7c, compare the blue lines on the left and right graphs). PRPF8 knockdown decreased spliceosomal binding at peaks 4/5 of both classes of BPs, and this led to stronger crosslinking at peaks 6/7 relative to peaks 4/5 at weak BPs, even though the peaks 4/5 are usually stronger. The signal at position B of weak BPs is almost completely lost upon PRPF8 knockdown, which likely reflects the absence of intron lariats due to perturbed splicing of introns with weak BPs (Fig. 7c). In conclusion, our results suggest that BP strength affects the assembly efficiency of spliceosomal factors at peaks 4/5, which could contribute to the variations between introns in their sensitivity to perturbed spliceosome function.

## Discussion

Here we established spliceosome iCLIP to study the interactions of endogenous snRNPs and accessory splicing factors on pre-mRNAs. We identified primary peaks of spliceosomal protein-pre-mRNA interactions, which precisely overlap with crosslinking profiles of 15 splicing factors. Moreover, the presence of lariat-derived reads in spliceosome iCLIP identified >50,000 BPs, which have canonical sequence and structural features. Due to the precise alignment of splicing factors to the positions of BPs, we could use their binding profiles to show that the assembly of U2 snRNP is primarily coordinated by the computationally predicted BPs, whilst the alternative BPs that are identified only by iCLIP or RNA-seq are more rarely used. Finally, we reveal the major effect of the position and strength of BPs on spliceosomal assembly, which can explain why distally located as well as weak BPs are particularly sensitive to perturbed spliceosome function upon PRPF8 KD. These findings demonstrate the broad utility of spliceosome iCLIP for simultaneous and transcriptome-wide analysis of the assembly of diverse spliceosomal components.

### The value of spliceosome iCLIP for identifying BPs

Experimental methods to identify BPs, which rely on reads from RNA-seq or iCLIP, are based on cDNAs derived from intron lariats. A caveat of these methods is that the stability of intron lariats depends on the kinetics of debranching and intron degradation, which may be affected by the properties of BPs. One study indicates that lariats formed at non-canonical BPs are less efficiently debranched^24^, which would increase the detection of non-canonical BPs by experimental methods. iCLIP captures a snapshot of RBP-RNA interactions that are in complex with spliceosome, which should minimize any biases of lariat stability. This could explain why the BPs identified by iCLIP contain a stronger consensus sequence and higher structural accessibility than the BPs that had been identified with lariat-spanning reads in RNA-seq. The reason for this difference may lie in the fact that lariats identified by iCLIP are in complex with the spliceosome at the time of crosslinking. The methods that rely on RNA-seq are expected to be more sensitive to the variable stability of intron lariats after their release from the spliceosome, which could lead to their greater propensity for detecting non-canonical BPs. The further value of spliceosome iCLIP is that, in addition to experiments under the medium condition, which serves for BP identification through lariat-derived cDNAs, experiments under the mild condition identify crosslinking of the RBPs in peaks 4/5 that align to BPs, thus enabling validation of BPs that is independent of variable lariat abundance (Fig. 5). Thus, a combined use of spliceosome iCLIP at both conditions is valuable to study the functionally relevant BPs, especially when combined with computational modelling of BPs^11^.

### The role of BP position and strength in spliceosomal assembly

We show that BP position and the computationally defined strength of BPs correlate with the relative binding of dozens of splicing factors around BPs. This is exemplified by strong binding of SF3 components at strong BPs, or BPs located close to 3’ss, whilst U2AF components bind stronger to weak BPs, or BPs located further from 3’ss. In cases of SF3B1, BUD13 and GPKOW, we observed enriched binding both at peaks 4/5 as well as 6/7, with reciprocal changes between the two peaks that depend on the features of BPs (Fig. 6 and 7). These RBPs are not known to bind at peaks 6/7, and it is plausible that signal at some peaks represents binding of U2AF or other spliceosomal factors that are co-purified during eCLIP. It is presently not possible to fully distinguish between direct and indirect binding, because the purified protein-RNA complexes have not been visualized after their separation on SDS-PAGE gels in eCLIP^12^. Nevertheless, our data clearly show that BP characteristics determine the balance of interactions between peaks 4/5 and 6/7 for a broad range of spliceosomal factors.

Our findings show a good convergence of transcriptomic insights with CryoEM studies of spliceosome structure. The RBPs with strongest enrichment at peaks 4/5 include SF3B4, SF3B1 and SF3A3, which are required for the ATP-dependent step of spliceosome assembly on the BPs^25^. This is in agreement with the ATP-dependence of peaks 4 and 5 *in vitro* and their disruption by PRPF8 KD. The binding positions of SF3B4 (peak at 26 nt upstream of BPs) and SF3A3 (peak at 15 nt upstream of BPs) is consistent with the structure of the human activated spliceosome, where SF3A3 (also referred to as SF3a60) binds to pre-mRNA at a position closer to the BP compared to SF3B4 (also referred to as SF3b49)^26^. Interestingly, while we observe binding peaks in the region 19-26 nt upstream of BPs in humans, the late spliceosomal components in yeast had their peak centered at ∼48-49 nt upstream of BPs^2^. In both cases, these contacts don’t overlap with any sequence motif, and thus their binding position appears to be defined by the assembly of the spliceosome on BPs. The constrained conformation of the larger spliceosomal complex appears to act as a molecular ruler that positions each associated RBP on pre-mRNAs at a specific distance from BPs.

In conclusion, spliceosome iCLIP monitors concerted pre-mRNA binding of many types of spliceosomal complexes with nucleotide resolution, allowing their simultaneous study due to the distinct position-dependent binding pattern of components that act at multiple stages of the splicing cycle. The method can be used to study endogenous spliceosome and BPs at multiple stages of development, and across tissues and species, without the need for protein tagging that was used in yeast^2,3^. Several spliceosomal components, including U2AF1, SF3B1 and PRPF8, are targets for mutations in myeloid neoplasms, retinitis pigmentosa and other diseases^27^. Spliceosome iCLIP could now be used to monitor global impacts of these mutations on spliceosome assembly in human cells. More generally, our study demonstrates the value of iCLIP for monitoring the position-dependent assembly and dynamics of multi-protein complexes on endogenous transcripts.

## Acknowledgements

We thank Livio Pellizzoni for the 18F6 monoclonal antibody, Miriam Llorian for help with the *in vitro* splicing reactions, Kathi Zarnack and Gregor Rot for help with the data analyses, and Lisa Strittmatter and members of Ule lab for helpful discussions and comments on the manuscript. This work was supported primarily by the European Research Council (206726-CLIP and 617837-Translate) and the Slovenian Research Agency (P2-0209, Z7-3665, J7-5460). CRS is supported by an Edmond Lily Safra fellowship. AMC is supported by a Wellcome Trust PhD Training Fellowship for Clinicians (110292/Z/15/Z). DP and VOW were supported by Medical Research Council programme grants MC_UU_12022/1 and MC_UU_12022/8 to ARV. The Francis Crick Institute receives its core funding from Cancer Research UK (FC001002), the UK Medical Research Council (FC001002), and the Wellcome Trust (FC001002).

## Author contributions

MB, CRS and JU conceived the project, designed the experiments and wrote the manuscript, with assistance of all co-authors. MB, CRS and ZW performed experiments, with assistance from JU, JK and CWS. NH performed most computational analyses, with assistance from CRS, TC, AMC and NML. VOW, DP and ARV provided crosslinked pellets from wild-type and PRPF8-depleted Cal51 cells.

## Supplementary legends

**Supplementary Fig. 1:**
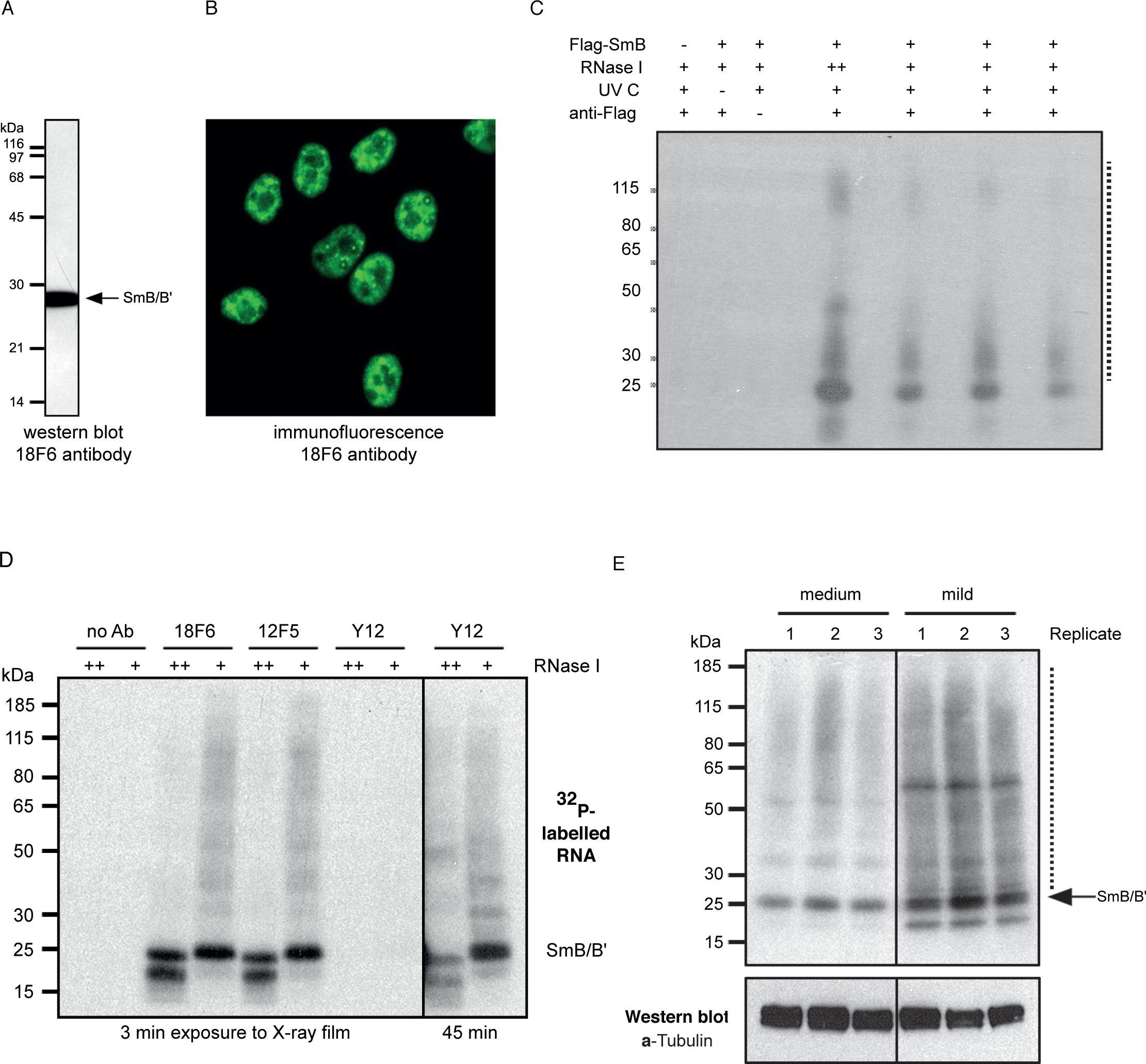
Quality control of spliceosome iCLIP with the anti-SmB/B’ antibodies. (a) Western blot analysis of total HeLa cell extract with 18F6 antibody reveals a single band of 28 kDa. (b) Analysis of HeLa cells by immunostaining with 18F6 and epifluorescence microscopy shows expected localization of SmB/B’ (a speckled nuclear pattern excluding nucleoli). (c) UV-crosslinked HEK FLP-in cells with stably integrated SmB-3×Flag were lysed under stringent conditions and subjected to partial RNase I digestion (+, final dilution 1:100,000; ++, final dilution 1:5,000). Spliceosomal RNPs were immunopurified with anti-Flag M2 antibody, RNA was 5’ end radiolabeled, and RNPs were subjected to denaturing gel electrophoresis and nitrocellulose transfer, an autoradiogram of which is shown. The interrupted line indicates the area on the nitrocellulose membrane cut out for purification of crosslinked RNP complexes. (d) Autoradiogram of crosslinked RNPs after immunopurification with the anti-SmB/B’ antibodies 18F6, 12F5 or Y12 (ab3138, Abcam). HeLa cell pellet was lysed in medium lysis buffer and subjected to high (++, final dilution 1:10,000) or low (+, final dilution 1:100,000) concentrations of RNase I. Lysates were split evenly between beads for immunopurification. RNAs of immunopurified RNP complexes were radiolabeled at the 5’ end followed by size-separation on denaturing gels and nitrocellulose transfer. The time below each panel indicates length of exposure during autoradiography. (e) UV-crosslinked mouse postnatal day 7 brains were lysed under medium or mild stringency conditions and subjected to partial RNase I digestion (final dilution 1:100,000). Spliceosomal RNPs were immunopurified with anti-SmB/B’ 18F6 antibody, RNA was 5’ end radiolabeled, and RNPs were subjected to denaturing gel electrophoresis and nitrocellulose transfer, an autoradiogram of which is shown in the upper panel. The interrupted line indicates the area on the nitrocellulose membrane cut out for purification of crosslinked RNP complexes. For Western blotting, the remainder of the supernatant following cell lysis and centrifugation was mixed with 4× NuPAGE LDS sample buffer (ThermoFisher) and equal sample volumes were separated by SDS-PAGE and transferred onto nitrocellulose membrane, which was incubated with anti-α-tubulin antibody (1:4,000, clone B-5-1-2, cat. no. T5168, Sigma-Aldrich).

**Supplementary Fig. 2:**
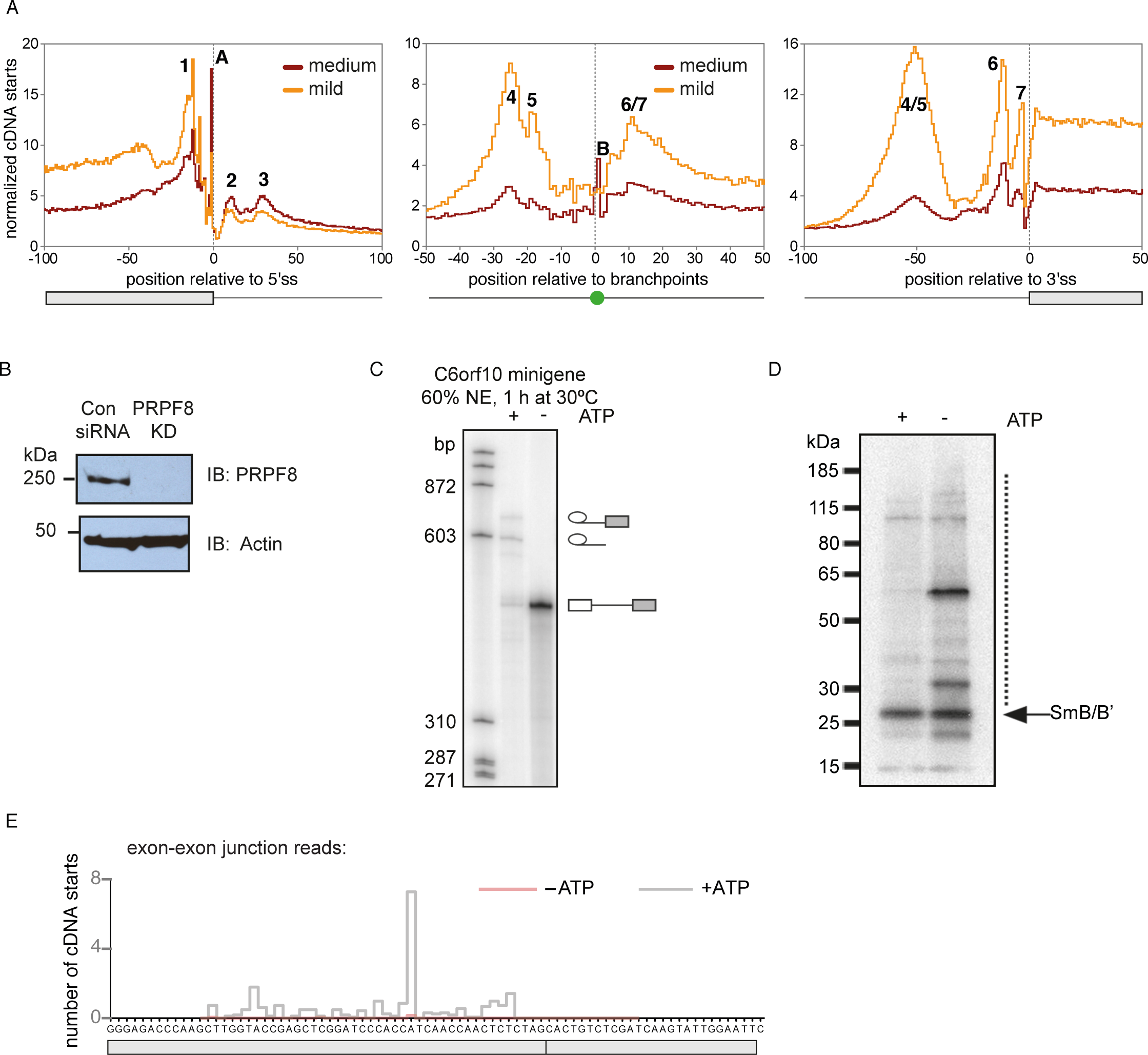
Analysis spliceosome iCLIP from cell extracts and *in vitro* splicing reactions. (a) RNA map of summarized crosslinking for spliceosome iCLIP performed under medium or mild conditions from mouse brain around the exon-intron, intron-exon junction and computationally top-scoring BP in each mouse intron^17^. (b) Immunoblot (IB) analysis of PRPF8 knockdown (KD) efficiency in Cal51 cells. (c) RNAs transcribed *in vitro* from a *C6orf10* minigene construct were incubated with HeLa nuclear extracts (NE) as part of *in vitro* splicing reactions in the presence or absence of ATP. Resulting splicing products and intermediates were resolved by denaturing gel electrophoresis and visualized by autoradiography. (d) *In vitro* splicing reactions were diluted in mild lysis buffer, subjected to low RNase I treatment (final dilution 1:200,000) and used for spliceosome iCLIP. Autoradiogram of crosslinked size-separated RNP complexes show the radiolabeled RNA that is crosslinked to RBPs. The interrupted line indicates the area cut out from the nitrocellulose membrane for extraction of crosslinked RNAs, which were used as a template for generating iCLIP cDNA libraries. (e) Normalized spliceosome iCLIP cDNA counts on the *C6orf10 in vitro* splicing product. Exons are marked by grey boxes. As expected, junction reads are almost exclusively present only in the +ATP library.

**Supplementary Fig. 3:**
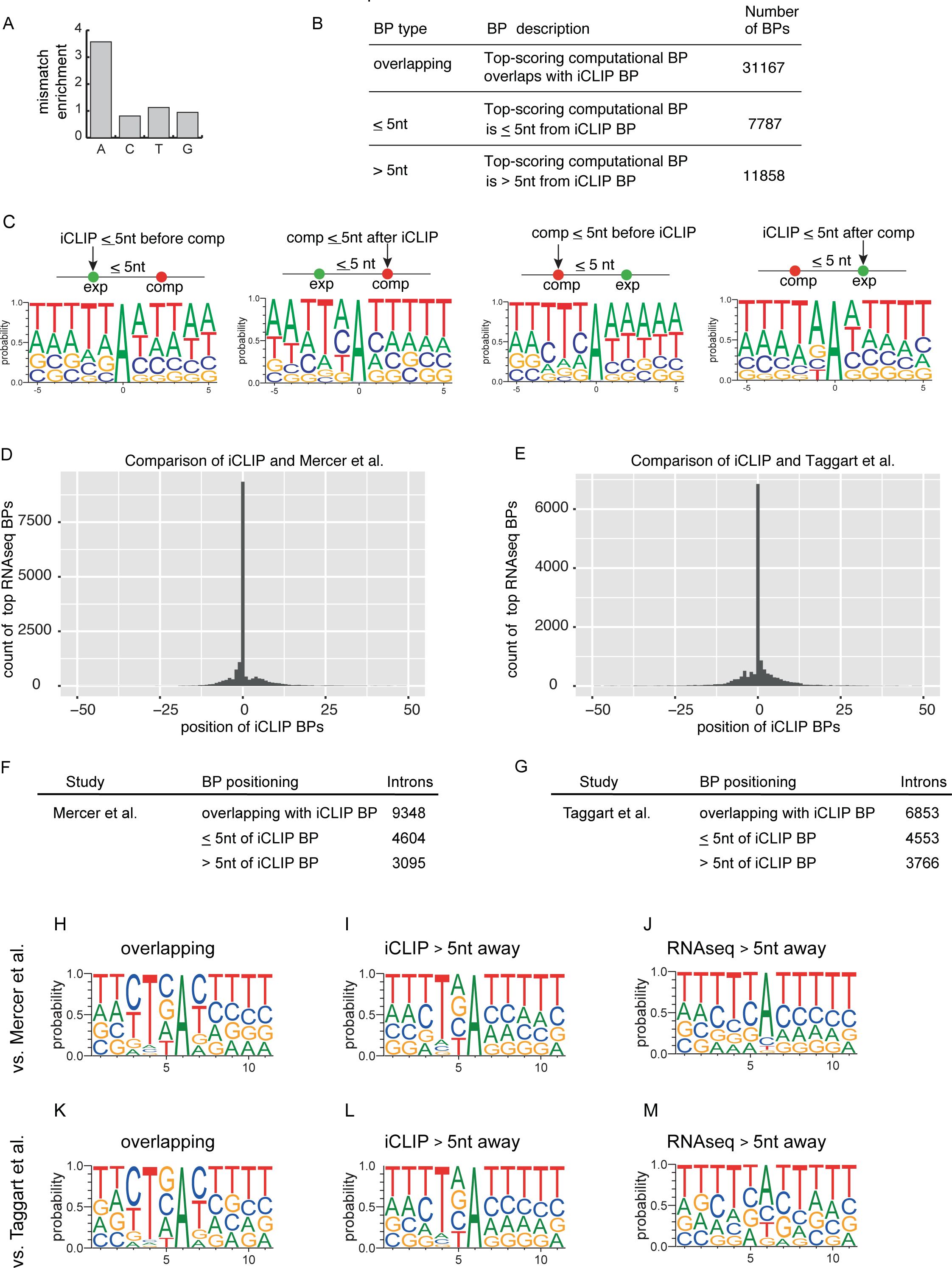
Comparison of BPs determined by spliceosome iCLIP to other methods. (a) Enrichment of mismatches at the first nucleotide of spliceosome iCLIP reads that overlap with ends of introns, compared to remaining iCLIP reads. (b) A table providing the number of BPs identified by spliceosome iCLIP (iCLIP BPs) in introns that also contain a computationally identified BP^11^. They are divided into three categories based on the distance between the iCLIP BP and the top-scoring computational BP in each intron. (c) Weblogo of four categories of non-overlapping BP that are ≤5 nt away from each other, centered either on iCLIP or computational BPs, and separated according to the relative position of iCLIP vs computational BP (upstream or downstream). (d) The distribution of top BPs identified by published RNA-seq^7^ (i.e., the BP with most lariat-spanning reads in each intron) around the BPs identified by spliceosome iCLIP (i.e., iCLIP BPs). (e) The distribution of top BPs identified by published RNA-seq^5^ (i.e., the BP with most lariat-spanning reads in each intron) around the BPs identified by spliceosome iCLIP (i.e., iCLIP BPs). (f) A table providing the number of BPs identified by spliceosome iCLIP (iCLIP BPs) in introns that also contain a BP assigned by lariat-spanning reads from RNA-seq^7^. They are divided into three categories based on the distance between the iCLIP BP and the top RNA-seq BP. (g) A table providing the number of BPs identified by spliceosome iCLIP (iCLIP BPs) in introns that also contain a BP assigned by lariat-spanning reads from RNA-seq^5^. They are divided into three categories based on the distance between the iCLIP BP and the top RNA-seq BP. (h) Weblogo of iCLIP BPs that overlap with RNA-seq BPs^7^. (i) Weblogo of iCLIP BPs that are >5 nt away from RNA-seq BP^7^. (j) Weblogo of RNA-seq BPs^7^ that are >5 nt away from iCLIP BP. (k) Weblogo of iCLIP BPs that overlap with RNA-seq BPs^5^. (l) Weblogo of iCLIP BPs that are >5 nt away from RNA-seq BP^5^. (m) Weblogo of RNA-seq BPs that are >5 nt away from iCLIP BP^5^.

**Supplementary Fig. 4:**
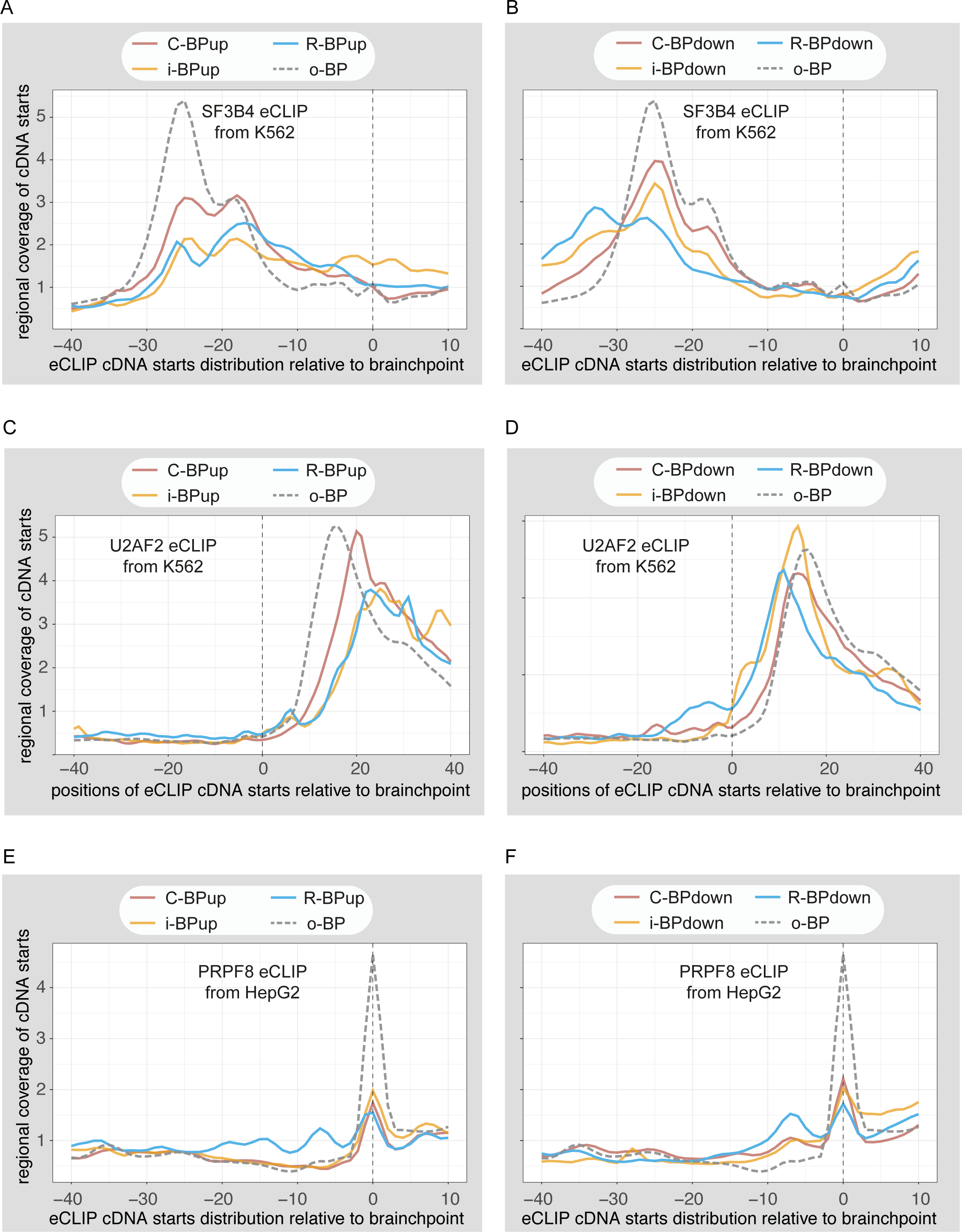
Spliceosome assembly at method-specific or overlapping BPs. RNA maps showing crosslinking (as cDNA starts from eCLIP experiments) of SF3B4 from K562 cells (a, b), of U2AF2 from K562 cells (c, d) and of PRPF8 from HepG2 cells (e, f) relative to BPs. BPs were categorized according to the method they were specifically detected by (spliceosome iCLIP, RNA-seq, computational prediction or overlapping) and in case of non-overlapping BPs, according to their location relative to each other: upstream (a, c, e) or downstream (b, d, f) of the other non-overlapping BP. Crosslinking of each RBP is regionally normalized to its average crosslinking over −100..50 nt region relative to 3’ss in order to most clearly allow comparisons between the relative positions of peaks for different RBPs.

**Supplementary Fig. 5:**
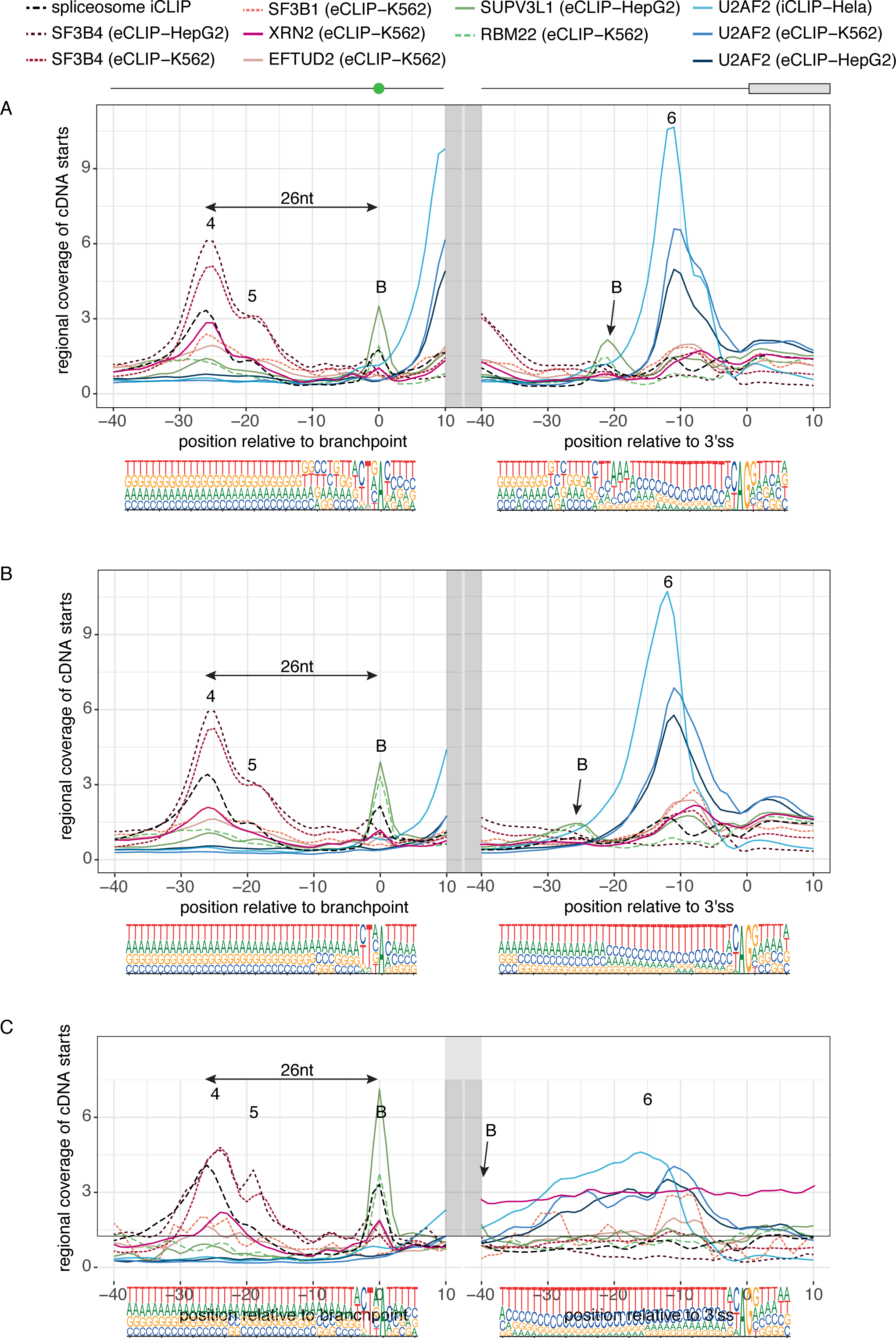
Crosslinking of many RBPs overlaps with peaks of spliceosomal crosslinking. (a) Crosslinking patterns of selected RBPs, as defined by cDNA starts of eCLIP or iCLIP in the indicated cell lines. Crosslinking of each is regionally normalized to its average crosslinking over −100..50 nt region relative to 3’ss in order to most clearly allow comparisons between the relative positions of peaks for different RBPs. All 3’ss that contain BPs within 17..23 nt upstream of the exon are chosen, and crosslinking is plotted in the region −40..10 nt relative to 3’ss, and −40..10 nt relative to BPs. (b) Same as (a), but for all 3’ss that contain BPs within 24..39 nt upstream of the exon. (c) Same as (a), but for all 3’ss that contain BPs within 40..65 nt upstream of the exon.

**Supplementary Fig. 6:**
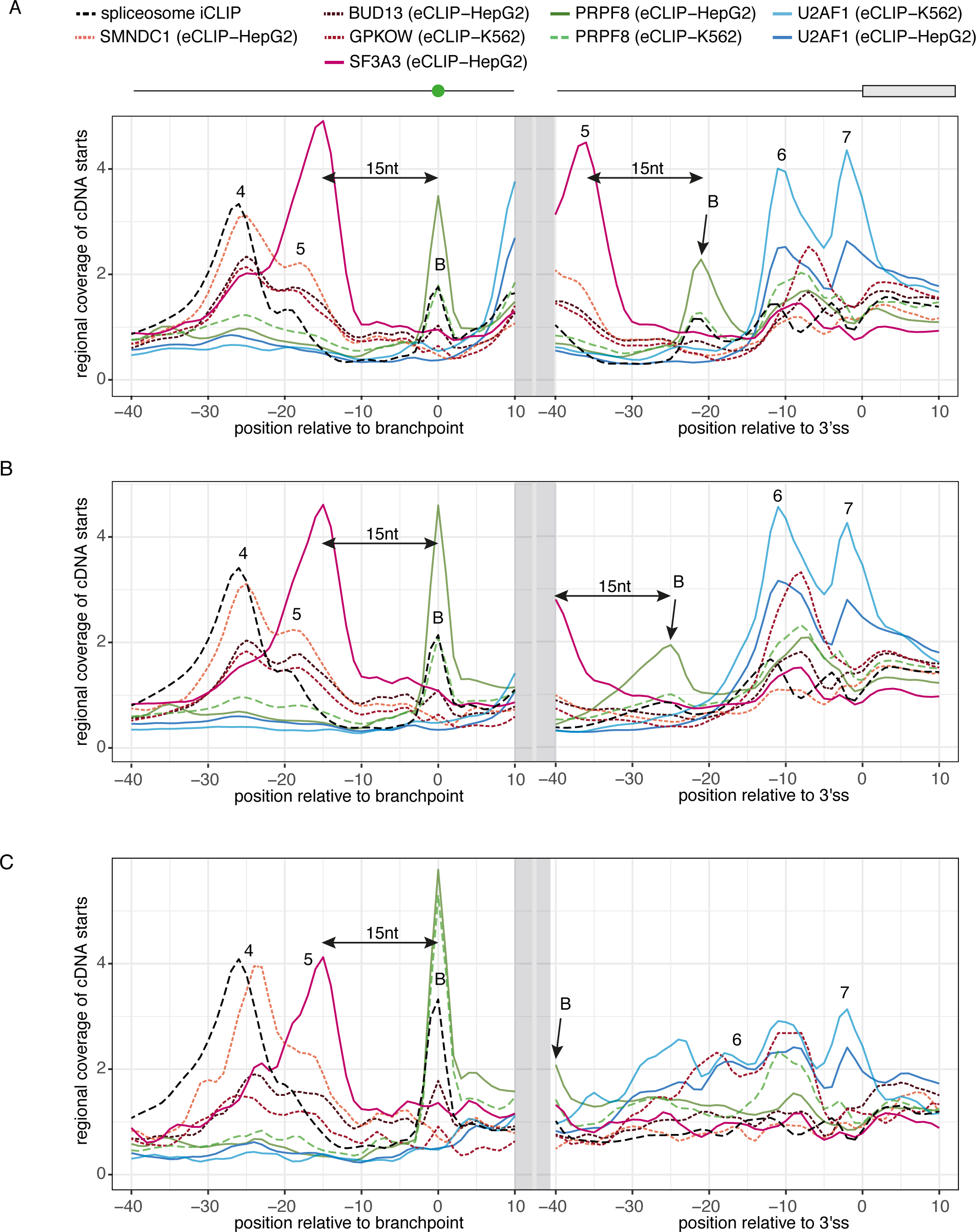
Crosslinking of many RBPs overlaps with peaks of spliceosomal crosslinking. (a) Crosslinking patterns of selected RBPs, as defined by cDNA starts of eCLIP or iCLIP in the indicated cell lines. Crosslinking of each is regionally normalized to its average crosslinking over −100..50 nt region relative to 3’ss in order to most clearly allow comparisons between the relative positions of peaks for different RBPs. All 3’ss that contain BPs within 17..23 nt upstream of the exon are chosen, and crosslinking is plotted in the region −40..10 nt relative to 3’ss, and −40..10 nt relative to BPs. (b) Same as (a), but for all 3’ss that contain BPs within 24..39 nt upstream of the exon. (c) Same as (a), but for all 3’ss that contain BPs within 40..65 nt upstream of the exon.

**Supplementary Fig. 7:**
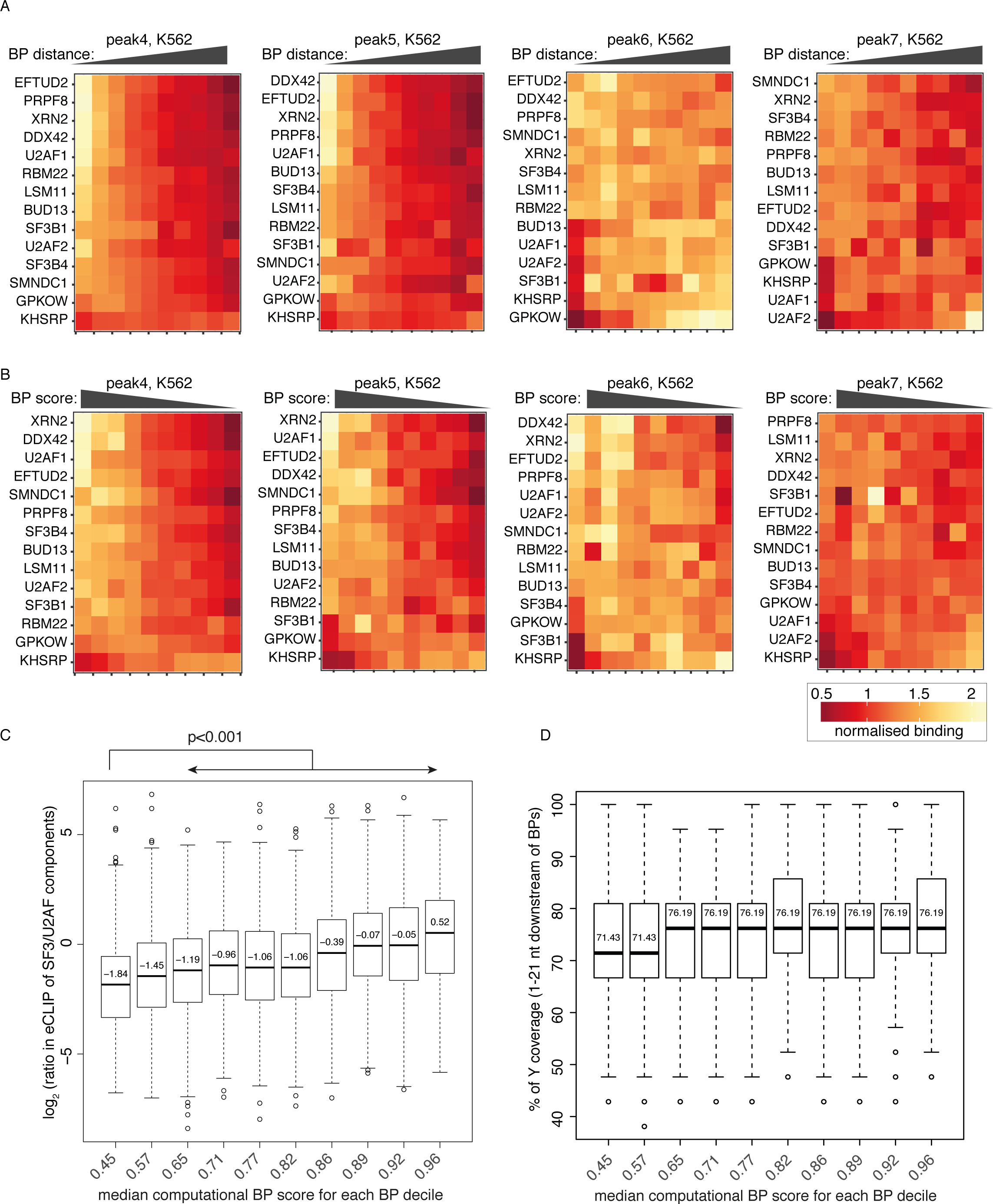
Relation of BP position and consensus score to binding of splicing factors. (b) Heatmaps depicting the normalized crosslinking of RBPs in peak regions around 10 groups of BPs that were categorized according to the distance of BPs from 3’ss. Crosslinks were derived as cDNA starts from eCLIP of K562 cells. (b) Heatmaps depicting the normalized crosslinking of RBPs in peak regions around 10 groups of BPs that were categorized according to the computational scores that define BP strength. Crosslinks were derived as cDNA starts from eCLIP of K562 cells. (c) BPs were divided into 10 quantiles based on their sequence consensus score, as determined previously^11^. The median score of each quantile is shown on the x-axis. The 4,410 BPs chosen for this analysis satisfied two criteria: 1) They were located 23-28 nt away from intron-exon junction, and 2) they contained a total of at least 30 crosslink events of SF3 (SF3B4-K562-eCLIP, SF3B4-HepG2-eCLIP and SF3A3-HepG2-eCLIP) in the region 35-10 nt upstream of BPs and U2AF (U2AF2-HepG2-eCLIP, U2AF2-K562-eCLIP and U2AF1-K562-eCLIP) in the region 5-25 nt downstream of BPs (the peak binding region of these RBPs). The y-axis shows the ratio in binding of SF3 relative to U2AF factors (data and positions as in the preceding sentence). P-values for the indicated comparisons were calculated by the pairwise Wilcoxon Rank Sum test. Box-plot elements are defined by center line, median; box limits, upper and lower quartiles; and whiskers, 1.5x interquartile range. (d) BPs were divided into 10 quantiles as in (c). The % of Ys (C or T) in the region 1-21 nt downstream of BPs is shown on the y-axis. Box-plot elements are defined by center line, median; box limits, upper and lower quartiles; and whiskers, 1.5x interquartile range.

## Methods

### Data and statistics

The spliceosome iCLIP data have been deposited on EBI ArrayExpress under the accession number E-MTAB-6950. These and published datasets referenced throughout this study are listed for convenience in Supplementary Table 7, including accession details. All statistical analyses were performed in the R software environment (version 3.1.3 and 3.3.2, https://www.r-project.org).

### Code availability

The code to identify BPs from spliceosome iCLIP reads is publicly available at the GitHub repository (https://github.com/nebo56/branch-point-detection-2).

### Preparation of Cal51 cells for iCLIP

Cal51 breast adenocarcinoma cells were prepared as described previously^4^. Briefly, cells were cultured in Dulbecco’s Modified Eagle Medium (DMEM, ThermoFisher) with 10% fetal calf serum (FCS, ThermoFisher) and 1× penicillin-streptomycin (P/S, ThermoFisher). For siRNA-mediated depletion of PRPF8, Cal51 cells were transfected with DharmaFECT1 (Dharmafect) with 25 nM siRNA targeting human *PRPF8*. Transfected cells were harvested 54 hrs later, exposed to UV-C light and used for iCLIP as described below. For collection of samples from different stages of the cell cycle, Cal51 cells were synchronized in G1/S by standard double thymidine block. Briefly, cells were treated with 1.5 mM thymidine for 8 hrs, washed and released for 8 hrs, then treated again with thymidine for a further 8 hrs. Cells were also collected 3 hrs (S-phase) and 7 hrs (G2) after release from the thymidine block.

### *In vitro* splicing

For *in vitro* splicing reactions, a *C6orf10* minigene construct containing exon 8 and 9 and 150 nt of the intron around both splice sites was produced (Fig. 2b). The minigene plasmid was linearized and transcribed *in vitro* using T7 polymerase with ^32^P-UTP. The transcribed RNA was then subjected to *in vitro* splicing reactions using HeLa nuclear extract. HeLa nuclear extract was depleted of endogenous ATP by pre-incubation and, for each reaction, 10 ng of RNA was incubated with 60% HeLa nuclear extract at 30°C with or without additional 0.5 mM ATP for 1 h in a 20 µl reaction. Afterwards, the reaction mixture was UV-crosslinked at 100 mJ/cm^2^ and stored at −80°C until further use. To visualize the splicing reaction products, proteinase K was added to the reaction mixture for 30 min at 37°C. The resulting RNA was phenol-extracted, precipitated and subjected to gel electrophoresis on a 5% polyacrylamide-urea gel.

### Spliceosome iCLIP protocol

For each experiment, three biological replicate samples of cDNA libraries were prepared (Supplementary Tables 2 and 3). The iCLIP method was done as previously described^10^, with the following modifications. Crosslinked cells or tissue were dissociated in the lysis buffer according to the stringency conditions (stringent, medium, mild; Supplementary Table 1) followed by sonication, low RNase I (AM2295, 100 U/µl, ThermoFisher) digestion and centrifugation. RNase at low concentration ensured that cDNAs are optimal size for comprehensive crosslink determination^14^. For denaturing, high-stringency experiment^10^, M2 anti-Flag antibody (Sigma) was used against the 3×Flag-SmB protein that had been stably integrated into HEK-293 FlpIn cells (Supplementary Fig. 1c). 6M Urea buffer was first used to lyse cell pellets, before being diluted down 1:9 with a Tween-20 containing IP buffer to allow for immuno-purification without denaturing of the M2 anti-Flag antibody, and then proceeded as described previously^14^.

Mouse brain tissue was used for initial experiments under mild and medium stringency conditions (Supplementary Fig. 1e), HeLa nuclear extract was used for *in vitro* splicing reactions (Supplementary Fig. 2c), and Cal51 cells were then used for all remaining experiments, since they have proven well-suited to understand the impact of spliceosomal perturbations on cell cycle^4^. For SmB/B’ immunopurification under medium and mild conditions, anti-SmB/B’ antibodies 12F5 (sc-130670, Santa Cruz Biotechnology) or 18F6 (as hybridoma supernatant, generated as described previously^9^) were used, which are different clones from the same immunization. These antibodies behave identically under immunopurification conditions (Supplementary Fig. 1d). For spliceosome iCLIP from mouse brain and *in vitro* splicing reactions, lysates were incubated with 50 μl monoclonal anti-SmB/B’ antibody 18F6, and for experiments from Cal51 cells, 12F5 anti-SmB/B’ antibody (Santa Cruz) was used. The antibody was pre-conjugated to 100 μl protein G Dynabeads (ThermoFisher) and rotated at 4°C followed by washing. As described previously, following immunopurification, RNA 3’ end dephosphorylation, ligation of the linker 5’-rAppAGATCGGAAGAGCGGTTCAG/ddC/-3’ to the 3’ end and 5’ end radiolabeling protein-RNA complexes were size-separated by SDS-PAGE and transferred onto nitrocellulose membrane. The regions corresponding to 28-180 kDa were excised from the membrane in order to isolate the bound RNA by proteinase K treatment. RNAs were reverse-transcribed in all experiments using SuperScript III reverse transcriptase at U/μl (ThermoFisher) and custom indexed primers (Table S2). Resulting cDNAs were subjected to electrophoresis on a 6% TBE-urea gel (ThermoFisher) for size selection. Purified cDNAs were circularized, linearized and amplified for high-throughput sequencing.

Identification of protein crosslink sites around splice sites, in particular at the peaks 4/5, was most efficient under the mild purification condition (Supplementary Fig. 2a). This condition was therefore used for analysis of spliceosomal assembly upon PRPF8 knockdown in Cal51 cells (Fig. 2a), and in the *in vitro* splicing reactions in HeLa nuclear extract (Fig. 2b). For the identification of BPs, we additionally used the medium condition, since it increases the frequency of cDNAs truncating at peak B (Supplementary Fig. 2a). For this purpose, spliceosome iCLIP was performed under medium purification conditions from Cal51 cells synchronized in G1, S and G2 phase. To maximise cDNA coverage, data from all synchronized cells was merged with the control Cal51 cells under mild condition for BP identification.

### Mapping of Sm iCLIP reads

We used mm9/NCBI37 and hg19/GRCh37 genome versions and Ensembl 75 gene annotation. Experimental and random barcode sequences of iCLIP sequenced reads were removed prior to mapping (Supplementary Table 2). We mapped the cDNAs to the genome with Bowtie 0.12.7 program using the parameters (-v 2 -m 1 -a --best --strata). The first 9 nt of the sequenced reads contain the experimental barcode to separate experimental replicates, and the random barcode, the latter of which allows to avoid artefacts caused by variable PCR amplification of different cDNAs^8^. We used these random barcodes to quantify the number of unique cDNAs at each genomic position by collapsing cDNAs with the same random barcode that mapped to the same starting position to a single cDNA. For analysis of crosslinking to snRNAs, we allowed sequences to map at up to 50 locations in the genome, but for all other analyses in the manuscript, we only allowed sequence mapping to a single location in the genome. For spliceosome iCLIP with the *C6orf10 in vitro* splicing substrate, sequence reads were first mapped to the unspliced substrates and the remaining reads were mapped to the spliced substrate allowing no mismatches. The nucleotide preceding the iCLIP cDNAs was used to define the crosslink sites.

### Mapping of eCLIP reads

For eCLIP sequencing data for all RBPs, we used GENCODE (GRCh38.p7) genome assembly and the STAR alignment (version 2.4.2a) using the following parameters from ENCODE pipeline: STAR --runThreadN 8 --runMode alignReads --genomeDir GRCh38 Gencode v25 --genomeLoad LoadAndKeep --readFilesIn read1, read2, --readFilesCommand zcat --outSAMunmapped Within –outFilterMultimapNmax 1 --outFilterMultimapScoreRange 1 --outSAMattributes All --outSAMtype BAM Unsorted – outFilterType BySJout --outFilterScoreMin 10 --alignEndsType EndToEnd --outFileNamePrefix outfile.

For the PCR duplicates removal, we used a python script ‘barcode collapse pe.py’ available on GitHub (https://github.com/YeoLab/gscripts/releases/tag/1.0), which is part of the ENCODE eCLIP pipeline (https://www.encodeproject.org/pipelines/ENCPL357ADL/).

### Normalization of crosslink positions for their visualization in the form of RNA maps

RNA maps were produced by summarizing the cDNA counts at each nucleotide using the previously developed RNA maps pipeline ^14,29^ relative to exon/intron and intron/exon boundaries and BPs on pre-mRNAs. The definition of intronic start and end positions was based on Ensembl version 75. Only introns longer than 300 nt were used to draw RNA maps in order to avoid detection of any RBPs that recognize 5’ss of introns.

In cases where we wished to compare the relative positions of crosslinking peaks between RBPs, we regionally normalized the summarized crosslinking of each RBP relative to the average crosslinking of the same RBP across the region 100 nt upstream and 50 nt downstream of the evaluated splice sites or branchpoints. Normalized values were then used to visualize the crosslinking in the form of RNA maps (Fig. 2, Supplementary Fig. 5 and 6).

To assess the role of BP characteristics on spliceosomal RBP assembly (Fig. 4, 6 and 7), we only examined the introns containing the 31,167 BPs that were identified both computationally and by iCLIP, which are likely the most reliable. We divided BPs into 10 categories based on BP position or score, and then normalized the summarized crosslinking of each RBP in each of the 10 BP categories relative to the average crosslinking of the same RBP across the region 100 nt upstream and 50 nt downstream of all the 31,167 evaluated BPs.

### Identification and comparison of branchpoints (BPs)

It has been shown that the spliceosomal C complexes harbor a salt-resistant RNP core containing U2, U5 and U6 snRNAs as well as the splicing intermediates including lariats that withstand treatment with 1M NaCl, whereas the spliceosomal B complexes were more likely dissociated under high-salt conditions^16^. This could explain why the medium purification condition is more suited than the mild condition to enrich for lariat cDNAs truncating at position B (Supplementary Fig. 2a). It is conceivable that the medium spliceosome iCLIP condition most strongly enriches spliceosomal C complexes, which are most effective for lariat detection. In contrast, the mild condition is expected to enrich additional B complexes that contain large amounts of SF3 components and have low proportion of lariats, in agreement with the strong enrichment of peaks 4 and 5 (Supplementary Fig. 2a). To identify the maximal diversity of BPs, we therefore pooled spliceosome iCLIP data produced under mild and medium purification conditions from Cal51 cells.

The first step to identify BPs used the spliceosome iCLIP reads that ended precisely at the ends of introns (we considered only introns that end in AG dinucleotide) after removal of the 3’ adapter. We noticed that these reads had an 3.5× increased frequency of mismatches on the A as the first nucleotide compared to remaining iCLIP reads (Supplementary Fig. 3a), indicating that these mismatches may have resulted from truncation at the three-way-junction formed at the BP (Fig. 2c). We therefore trimmed the first nucleotide from the read if it contained a mismatch at the first position that corresponded to a genomic adenosine. We then used spliceosome iCLIP from Cal51 cells to identify all reads that ended precisely at the ends of introns and defined the position where these reads started and assessed the random barcode nucleotides that are present at the beginning of each iCLIP read to count the number of unique cDNAs at each position. The nucleotide preceding the read start corresponds to the position where cDNAs truncated during the reverse transcription, and we selected the genomic A that had the highest number of truncated cDNAs as the candidate BP. If two positions with equal number of cDNAs were found, we selected the one closer to the 3’ss. For all branchpoint analyses, we only assessed protein-coding genes with FPKM>10 in the RNA-seq data, which identified 35,056 BP positions.

In the second step of analysis, we considered all cDNAs (regardless of where they ended), but including trimming of the first nucleotide if there was a mismatch with the genomic A. We then overlapped cDNA truncation sites with computationally predicted BPs in the last 100 nt of intron^17^. If this analysis identified a position with a higher cDNA count than the initial analysis (or if the initial analysis didn’t identify any BP in the same intron), then the newly identified position was assigned as the BP. For introns where no BP was identified by either the first or second step in the analysis, we assessed computationally predicted BPs located further than 100 nt from the 3’ss, and if any of these overlapped with a truncating cDNA, we assigned the position closest to the 3’ss as the BP. Together, this identified 50,812 BPs in genes with FPKM>10. The coordinates of these BPs were used for analyses presented in the Figures 4-7. We additionally identified 13,496 BPs in introns of lowly expressed genes, but these were not used for any further analyses.

We also attempted to use truncated cDNAs from PRPF8 eCLIP for discovery of BPs, but found that the number of cDNAs overlapping with intron ends was much smaller than in spliceosome iCLIP, and was insufficient for BP discovery. This is most likely because the high amount of non-specific background signal in PRPF8 eCLIP, which leads to a lower proportion of cDNAs that align to the BPs.

Bedtools Intersect command using option –u was used to compare BP coordinates from spliceosome iCLIP to the BPs identified in previous studies. We restricted this comparison to introns where BPs were detected by all three datasets (iCLIP, RNA-seq and computational prediction).

To define a single ‘computational BP’ per intron, the BP positions computationally predicted for each intron in hg19 were obtained from11: http://bejerano.stanford.edu/labranchor/, and top scoring BP in each intron was use. To define a single ‘RNA-seq BP’ per intron, we used the BP with most lariat-spanning reads in each intron.

### Analysis of pairing probability

Computational predictions of the secondary structure were performed by RNAfold function from Vienna Package (https://www.tbi.univie.ac.at/RNA/) with default parameters^28^. The RNAfold results are provided in a customized format, where brackets are representing the double stranded region on the RNA and dots are used for unpaired nucleotides. We measured the density of pairing probability by summing the paired positions into a single vector.

### Identification of RBPs overlapping with spliceosomal peaks

For RBP enrichment in Fig. 4, we used the eCLIP data from the ENCODE consortium^15^, together with available iCLIP experiments from our lab (which are all listed in^22^), to see if any of the proteins are enriched in the region of spliceosomal peaks. In total this included 157 eCLIP samples of 68 RBPs in the HepG2 cell line, and 89 RBPs in the K562 cell, and iCLIP samples of 18 RBPs from different cell lines (Supplementary Table 5). Next, we intersected cDNA-starts from each sample to the −100 to +50 nt region relative to the 3’ss and used it as control for each of the following peaks: Peak 4 (−23 nt..−29 nt relative to BP), Peak 5 (−21 nt..−17 nt relative to BP), Peak B (−1 nt..1 nt relative to BP), Peak A (−1 nt..1 nt relative to 5’ss), Peak 6 (−11 nt..−10 nt relative to 3’ss), Peak 7 (−3 nt..−2 nt relative to 3’ss). The positions of these peaks were determined based on crosslink enrichments in spliceosome iCLIP.

## References

1. Fica, S. M. & Nagai, K. Cryo-electron microscopy snapshots of the spliceosome: structural insights into a dynamic ribonucleoprotein machine. Nat Struct Mol Biol 24, 791-799, doi:10.1038/nsmb.3463 (2017).

2. Chen, W. et al. Transcriptome-wide Interrogation of the Functional Intronome by Spliceosome Profiling. Cell 173, 1031-1044 e1013, doi:10.1016/j.cell.2018.03.062 (2018).

3. Burke, J. E. et al. Spliceosome Profiling Visualizes Operations of a Dynamic RNP at Nucleotide Resolution. Cell 173, 1014-1030 e1017, doi:10.1016/j.cell.2018.03.020 (2018).

4. Wickramasinghe, V. O. et al. Regulation of constitutive and alternative mRNA splicing across the human transcriptome by PRPF8 is determined by 5’ splice site strength. Genome Biol 16, 201, doi:10.1186/s13059-015-0749-3 (2015).

5. Taggart, A. J. et al. Large-scale analysis of branchpoint usage across species and cell lines. Genome Res 27, 639-649, doi:10.1101/gr.202820.115 (2017).

6. Pineda, J. M. B. & Bradley, R. K. Most human introns are recognized via multiple and tissue-specific branchpoints. Genes Dev 32, 577-591, doi:10.1101/gad.312058.118 (2018).

7. Mercer, T. R. et al. Genome-wide discovery of human splicing branchpoints. Genome Res 25, 290-303, doi:10.1101/gr.182899.114 (2015).

8. König, J. et al. iCLIP reveals the function of hnRNP particles in splicing at individual nucleotide resolution. Nat Struct Mol Biol 17, 909-915, doi:nsmb.1838 [pii] 10.1038/nsmb.1838 (2010).

9. Carissimi, C., Saieva, L., Gabanella, F. & Pellizzoni, L. Gemin8 is required for the architecture and function of the survival motor neuron complex. J Biol Chem 281, 37009-37016, doi:M607505200 [pii] 10.1074/jbc.M607505200 (2006).

10. Huppertz, I. et al. iCLIP: protein-RNA interactions at nucleotide resolution. Methods 65, 274-287, doi:10.1016/j.ymeth.2013.10.011 (2014).

11. Paggi, J. M. & Bejerano, G. A sequence-based, deep learning model accurately predicts RNA splicing branchpoints. RNA 24, 1647-1658, doi:10.1261/rna.066290.118 (2018).

12. Lee, F. C. Y. & Ule, J. Advances in CLIP Technologies for Studies of Protein-RNA Interactions. Mol Cell 69, 354-369, doi:10.1016/j.molcel.2018.01.005 (2018).

13. Sugimoto, Y. et al. Analysis of CLIP and iCLIP methods for nucleotide-resolution studies of protein-RNA interactions. Genome biology 13, R67, doi:10.1186/gb-2012-13-8-r67 (2012).

14. Haberman, N. et al. Insights into the design and interpretation of iCLIP experiments. Genome Biol 18, 7, doi:10.1186/s13059-016-1130-x (2017).

15. Van Nostrand, E. L. et al. A Large-Scale Binding and Functional Map of Human RNA Binding Proteins. bioRxiv, doi:10.1101/179648 (2017).

16. Bessonov, S., Anokhina, M., Will, C. L., Urlaub, H. & Luhrmann, R. Isolation of an active step I spliceosome and composition of its RNP core. Nature 452, 846-850, doi:10.1038/nature06842 (2008).

17. Corvelo, A., Hallegger, M., Smith, C. W. & Eyras, E. Genome-wide association between branch point properties and alternative splicing. PLoS computational biology 6, e1001016, doi:10.1371/journal.pcbi.1001016 (2010).

18. Wang, Z. et al. iCLIP predicts the dual splicing effects of TIA-RNA interactions. PLoS Biol 8, e1000530, doi:10.1371/journal.pbio.1000530 (2010).

19. Tollervey, J. R. et al. Characterizing the RNA targets and position-dependent splicing regulation by TDP-43. Nat Neurosci 14, 452-458, doi:nn.2778 [pii] 10.1038/nn.2778 (2011).

20. Rogelj, B. et al. Widespread binding of FUS along nascent RNA regulates alternative splicing in the brain. Sci Rep 2, 603, doi:10.1038/srep00603 (2012).

21. Zarnack, K. et al. Direct Competition between hnRNP C and U2AF65 Protects the Transcriptome from the Exonization of Alu Elements. Cell 152, 453-466, doi:10.1016/j.cell.2012.12.023 (2013).

22. Attig, J. et al. Heteromeric RNP Assembly at LINEs Controls Lineage-Specific RNA Processing. Cell 174, 1067-1081 e1017, doi:10.1016/j.cell.2018.07.001 (2018).

23. Gozani, O., Feld, R. & Reed, R. Evidence that sequence-independent binding of highly conserved U2 snRNP proteins upstream of the branch site is required for assembly of spliceosomal complex A. Genes Dev 10, 233–243 (1996).

24. Hartmuth, K. & Barta, A. Unusual branch point selection in processing of human growth hormone pre-mRNA. Mol Cell Biol 8, 2011–2020 (1988).

25. Wahl, M. C., Will, C. L. & Lührmann, R. The spliceosome: design principles of a dynamic RNP machine. Cell 136, 701-718, doi:S0092-8674(09)00146-9 [pii] 10.1016/j.cell.2009.02.009 (2009).

26. Zhang, X. et al. Structure of the human activated spliceosome in three conformational states. Cell research 28, 307-322, doi:10.1038/cr.2018.14 (2018).

27. Scotti, M. M. & Swanson, M. S. RNA mis-splicing in disease. Nat Rev Genet 17, 19-32, doi:10.1038/nrg.2015.3 (2016).

28. Lorenz, R. et al. ViennaRNA Package 2.0. Algorithms for molecular biology : AMB 6, 26, doi:10.1186/1748-7188-6-26 (2011).

29. Chakrabarti, A., Haberman, N., Praznik, A., Luscombe, N. M. & Ule, J. Data Science Issues in Studying Protein–RNA Interactions with CLIP Technologies. Annual Review of Biomedical Data Science Vol. 1, doi:https://doi.org/10.1146/annurev-biodatasci-080917-013525 (2018).

